# Functional tunability from a distance: Rheostat positions influence allosteric coupling between two distant binding sites

**DOI:** 10.1101/632422

**Authors:** Tiffany Wu, Liskin Swint-Kruse, Aron W. Fenton

## Abstract

For protein mutagenesis, a common expectation is that important positions will behave like on/off “toggle” switches (*i.e.*, a few substitutions act like wildtype, most abolish function). However, there exists another class of important positions that manifests a wide range of functional outcomes upon substitution: “rheostat” positions. Previously, we evaluated rheostat positions located near the allosteric binding sites for inhibitor alanine (Ala) and activator fructose-1,6-bisphosphate (Fru-1,6-BP) in human liver pyruvate kinase. When substituted with multiple amino acids, many positions demonstrated moderate rheostat effects on allosteric coupling between effector binding and phosphoenolpyruvate (PEP) binding in the active site. Nonetheless, the combined outcomes of all positions sampled the full range of possible allosteric coupling (full tunability). However, that study only evaluated allosteric tunability of “local” positions, *i.e*., positions were located near the binding sites of the allosteric ligand being assessed. Here, we evaluated tunability of allosteric coupling when mutated sites were distant from the allosterically-coupled binding sites. Positions near the Ala binding site had rheostat outcomes on allosteric coupling between Fru-1,6-BP and PEP binding. In contrast, positions in the Fru-1,6-BP site exhibited modest effects on coupling between Ala and PEP binding. Analyzed in aggregate, both PEP/Ala and PEP/Fru-1,6-BP coupling were again fully tunable by amino acid substitutions at this limited set of distant positions. Furthermore, some positions exhibited rheostatic control over multiple parameters and others exhibited rheostatic effects on one parameter and toggle control over a second. These findings highlight challenges in efforts to both predict/interpret mutational outcomes and engineer functions into proteins.

## Introduction

In mutagenesis studies to evaluate the contributions of important amino acid positions to protein function, our collective experience has led to a common expectation: For a given position, a few substitutions result in wildtype function, but most substitutions abolish function (*i.e.*, function is “toggled” on or off like a light switch)(Figure 1). However, Gray et al^1^ showed that most substitution studies on which our expectations are based have focused on conserved positions, which biases our collective experience. Recently, we selected a broader range of positions for mutagenesis and in doing so, we identified another type of mutagenesis outcome: When these positions were substituted with various amino acids, a wide range of functional outcomes was obtained. When rank-ordered, the outcomes showed a continuum of functional magnitudes, analogous to electronic rheostats (*e.g.*, a dimmer switch)^2^. We further found that a histogram analyses can be used to quantify the degree of rheostat character^3^.

**Figure 1.**
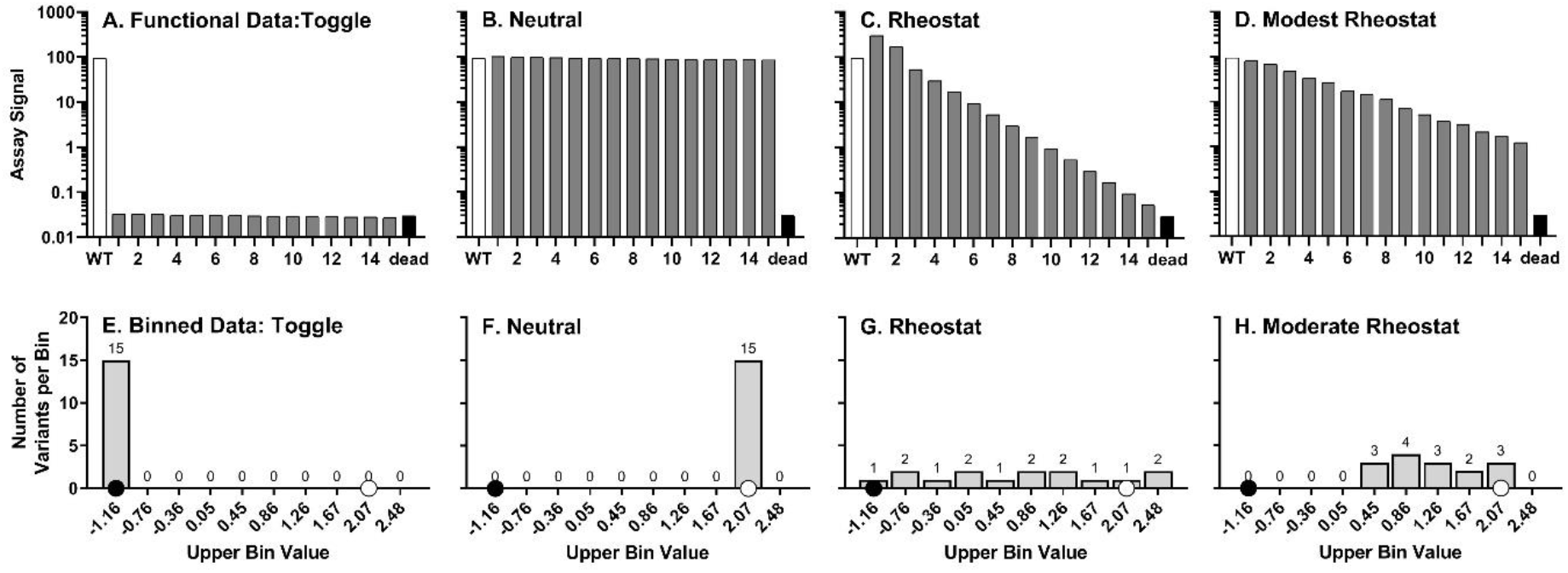
Simulated examples of neutral, toggle, rheostat, and modest rheostat positions. On the x-axis in panels A-D, numbers are used as placeholders for amino acid substitution type. Panel A shows a toggle position for which all substitutions abolish function: this simulated example gives rise to the maximum possible toggle score. For prior experimental data ^14^, toggle positions were defined when at least 64% of substitutions abolished function (*i.e.*, “dead”) and at most 16% of the substitutions exhibited wildtype function. Panel B shows a neutral position, for which most substitutions are like wildtype. Panel C shows a rheostat position for which substitution values range from wildtype to “dead”. In our experimental identification of rheostat positions, we have also found examples that do not perfectly conform to the rheostat profile but nevertheless exhibit rheostat-like properties^3^. One such example is shown in panel D. The key features for quantifying the degree of a “rheostat” position include the total range of possible outcomes, which are usually delimited by the parameter values of wildtype and “dead” protein^3^. For subsequent histogram analyses, this range is first divided into bins; here, the histograms in panels E-H correspond to the simulated data in panels A-D. The values on the x-axis indicate the upper-bound value of a bin in which all variants possess logarithmic transformations of functional value lesser than or equal to the label of the bin and neither lesser nor equal to the next smaller bin. A white dot is used to indicate the bin that includes wildtype data and a black dot is used to indicate the bin that corresponds to “dead” (*e.g.*, “no allostery”) function.

Once rheostat positions were recognized to exist, we initiated studies to evaluate the occurrence of rheostat positions across a range of proteins with different, quantifiable functions^3^. One such protein was human liver pyruvate kinase (hLPYK), a model system for allosteric regulation. The affinity of hLPYK for its substrate in the active site is regulated by two different allosteric effectors. Thus, five functional parameters are used to characterize (i) enhancement of substrate (phosphoenolpyruvate; PEP) affinity by the allosteric activator (fructose-1,6-bisphosphate; Fru-1,6-BP) and (ii) reduction of PEP affinity caused by the allosteric inhibitor (alanine; Ala) (Table 1)^4–6^. When scored for their rheostat character^3^, most of the individual positions in hLPYK allosteric sites had modest rheostat scores for either binding of the allosteric ligand or allosteric coupling between effector and substrate binding.

**Table 1:** Parameter Definition

| Table 1: Parameter Definition |  |
| --- | --- |
| $K_{a-PEP}$ | PEP apparent affinity |
| $K_{ix-Ala}$ | Ala binding |
| $K_{ix-FBP}$ | Fru-1,6-BP binding |
| $Q_{ax-Ala}$ | Allosteric coupling between<br>PEP binding and Ala<br>binding |
| $Q_{ax-FBP}$ | Allosteric coupling<br>between PEP and Fru-1,6-BP binding |

In contrast to the evaluation of single positions, when outcomes for all mutated positions from one allosteric site were combined into a single histogram, high rheostat scores were calculated for both allosteric coupling parameters (Figure 2). This led us to refine our definitions: A “rheostat position” is a single position for which substitutions achieve a full range of functional outcomes between the absence of function (“dead”) and “better than wildtype”. “Full tunability” describes the scenario in which a complete range of functional outcomes can be obtained by substituting a subset of positions. In histogram analyses, tunability increases as more bins have at least one entry (Figure 2). Note that full tunability (all bins having at least one entry) can be obtained even if the individual component positions are modestly or poorly rheostatic.

**Figure 2.**
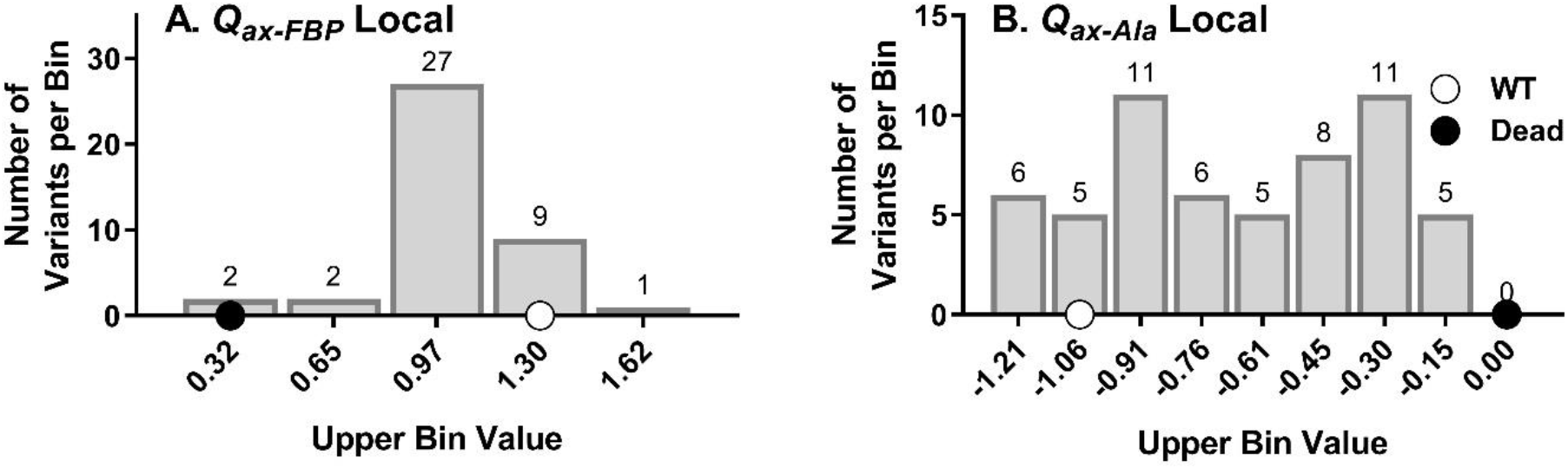
Binning of allosteric coupling functions for mutations at “local” positions; functional data were taken from the previous study^3^ but analyzed as described in this paper for consistency. All substitutions at seven positions in the Fru-1,6-BP binding site were evaluated for their effects on PEP/Fru-1,6-BP allosteric coupling (*Q*_*ax-FBP*_), and all substitutions at eight positions in the Ala binding site were evaluated for their effects on PEP/Ala allosteric coupling (*Q*_*ax-Ala*_). Data shown here are a compilation of the measurements of allosteric coupling obtained for all substitutions at all positions in each of the allosteric binding sites. “Tunability” of these parameters is apparent as a high percentage of bins of logarithmic transformations of functional data have at least one entry (see numbers above each bin). In other words, at least one substitution in each binding site results in one of the possible functional outcomes (histogram bins) that spans the range between no allostery/”dead” (black dot) and wildtype allosteric coupling (white dot).

Our previous study identified full tunability for “local” allosteric binding sites. That is, mutations were made in the allosteric effector binding site involved in the allosteric coupling parameter studied (*e.g.*, positions in the Fru-1,6-BP binding site were evaluated for effects on PEP/Fru-1,6-BP allosteric coupling). Therefore, we were curious whether allosteric coupling could be fully tunable when substituted positions were distant from the two allosterically-coupled binding sites. In hLPYK, this could be assessed by considering whether substitutions in the Fru-1,6-BP binding site altered coupling between PEP and Ala, and *vice versa*. Here, we show tunability was again high from a small set of positions even though those positions were distant from the two allosteric sites being studied. Several positions exhibited rheostat control over multiple parameters. Furthermore, two positions simultaneously acted as a rheostat for one parameter and a toggle for a second parameter, highlighting the challenges that arise when predicting/interpreting mutational outcomes and rationally engineering functional variation into existing proteins.

## Materials and Methods

All methods were performed in accordance with relevant guidelines. In particular, all use of recombinant DNA was according to the NIH Guidelines and was approved by the KUMC Institutional Biosafety Committee.

All hLPYK mutations were previously created^7,8^. This group of mutations include substitutions at seven positions in the Fru-1,6-BP binding site and eight positions in the Ala binding site. At each position, 9-16 substitutions were created for a total of 197 variants of hLPYK. Protein preparation, assays of enzymatic activity, and data analysis were carried out as previously reported. In short, using substrate titrations of enzymatic activity, the apparent affinity of hLPYK for its substrate, PEP, was evaluated over a concentration range for either Fru-1,6-BP or Ala. For each allosteric ligand, the coupling constant *Q*_*ax*_ was calculated to characterize allosteric function ^4,9–11^:

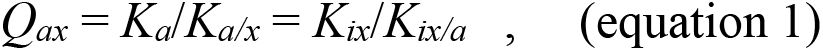

where *K*_*a*_ is the apparent affinity for ligand A (*e.g.*, substrate *K*_*a-PEP*_) in the absence of ligand X (*e.g.*, effector Ala or Fru-1,6-BP); *K*_*a/x*_ is the apparent affinity for ligand A in the presence of saturating ligand X; *K*_*ix*_ is the affinity for ligand X (*e.g.*, *K*_*ix-Ala*_ and *K*_*ix-FBP*_) in the absence of ligand A; and *K*_*ix/a*_ is the affinity of the protein for ligand X in the presence of saturating ligand A (Table 1).

When a variant failed to demonstrate detectable activity, protein preparation was replicated at least three times before designating the protein as catalytically non-active. No additional steps were taken to concentrate protein or assay immediately after cell lysis, both of which are approaches that can better detect enzymatic activity in a destabilized mutant protein. Additionally, no further evaluations of protein properties were performed for non-active mutant proteins.

Once affinity, binding, and allosteric coupling parameters were determined for variants at each position, the degree of rheostat character was evaluated using the RheoScale calculator^3^. For each position, this calculator uses all available variants to quantify its rheostat, toggle, and neutral character *via* histogram analyses. Rheostat scores describe how well the functional data sample the histogram bins. Toggle scores reflect the number of variants that fall into the same bin as “dead” protein. Neutral scores reflect the number of variants that fall into a histogram bin centered on the value of the wildtype protein; these calculations are extensively discussed in ^12^.

To identify the appropriate width and number of histogram bins, the RheoScale calculator assessed the full experimental dataset for (i)minimal and maximal values for the parameter being assessed, (ii) average error of experimental measurements, and (iii) average number of variants available per position. Since completing our first study of “local” hLPYK rheostat positions^3^, we have obtained and collated a much larger dataset of hLPYK parameters and used them to refine RheoScale histogram analyses ^12,13^. These include measurements for additional variants and multiple, independent measurements of wildtype hLPYK (Supplemental Information). The latter was used to estimate experimental error for all variants in this study. (Prior RheoScale analyses used errors of the fit which are smaller than experimental errors.) Revised “local” scores are reported in the supplement and used for comparison to the current “distant” scores. Note that neutral scores were calculated using the revised methods of ^12^ for both local and distant scores.

For most of the RheoScale calculations, “local” rheostat scores showed little change. However, for *Q*_*ax-FBP*_, two factors became apparent. First, replicates of wildtype hLPYK showed that this parameter had larger experimental error than previously estimated (and larger than other parameters). This decreased the number of bins used in histogram analyses to 5 when the histogram range was delimited by wildtype and “dead” (zero allosteric coupling). Second, several additional variants had “better” *Q*_*ax-FBP*_ values than wildtype hLPYK; this expanded the maximum range of the histogram. Some of these variants had such large effects on *Q*_*ax-FBP*_, and thus on the maximal range, that if delimited by maximum score and zero, all other variants would be compressed into a small percent of bins. Empirically, we found that this under-represented the rheostat character of positions for which substitutions accessed the full range between wildtype and “dead” allosteric coupling. Thus, we chose to classify all “better than wildtype” values into a single bin and reduce the number of bins between wildtype and “dead” by 1 from the RheoScale-recommended bin number. In doing so, the number of bins used in histogram analysis remains the same as the recommended bin number. Furthermore, the bin containing wildtype was centered on the average of all wildtype measurements. This methodology of n-1.5 bins (where n is the recommended number of bins) in the wildtype average to “dead” range and one “better than wildtype” bin was used for all parameters.

## Results

In this work, we evaluated seven positions in/near the Fru-1,6-BP site^8^ and eight positions in/near the Ala binding site^7^. Nine to 16 substitutions were introduced at each of these positions for a total of 197 substitutions. We previously reported *K*_*a-PEP*_, *K*_*ix-Ala*_, and *Q*_*ax-Ala*_ for variants in the Ala binding site and *K*_*a-PEP*_, *K*_*ix-FBP*_, and *Q*_*ax-FBP*_ for variants in the Fru-1,6-BP binding site^3^. For this work, we collected an additional 18,000 assays to determine *K*_*ix-FBP*_ and *Q*_*ax-FBP*_ for variants in the Ala binding site and *K*_*ix-Ala*_ and *Q*_*ax-Ala*_ for variants in the Fru-1,6-BP binding site. As previously noted^3^, most substitutions at position 483 resulted in a catalytically inactive protein. Since this prevented further evaluation of the binding of effectors and allosteric coupling, this position was not considered further.

To quantify the composite functional results for the 16 positions of this study, we used the RheoScale calculator to generate three scores: a rheostat score, a toggle score, and a neutral score. All three scores range from 0 to 1, with 1 being the perfect manifestation of the substitution character (*e.g*., a perfect rheostat position). Empirically, for datasets the size of the hLPYK set, we have found that rheostat scores above 0.5 indicate strong rheostat character of the position^3^. In contrast to rheostat scores, neutral scores must be above 0.7 to be considered a high score ^12^. For toggle scores, we previously defined a strong toggle threshold of “at least 8 severe variants” ^14^. In example positions with 12-13 substitutions, this definition effectively results in a RheoScale toggle score of 0.64 or greater. As detailed in Methods, these and other new hLPYK data were used to refine RheoScale parameters; thus previously-published data for “local” substitutions were reanalyzed with new parameters to maintain consistency in comparisons. The revised scores are reported in the supplement, as are all experimental measurements for the “distant” substitutions and histograms for individual positions.

Analyses of the distant experimental measurements are shown in Figure 3. On the left-hand side of each panel, the influence of positions in the Ala binding site on Fru-1,6-BP binding and PEP/Fru-1,6-BP allosteric coupling is shown. Overall, rheostat scores for *Q*_*ax-FBP*_ (most above 0.5) were notably larger than *K*_*ix-FBP*_. None of the parameters showed neutral scores above the 0.7 neutral score threshold. Alltoggle scores for mutants in the Ala binding site were below the 0.64 toggle score cutoff.

**Figure 3.**
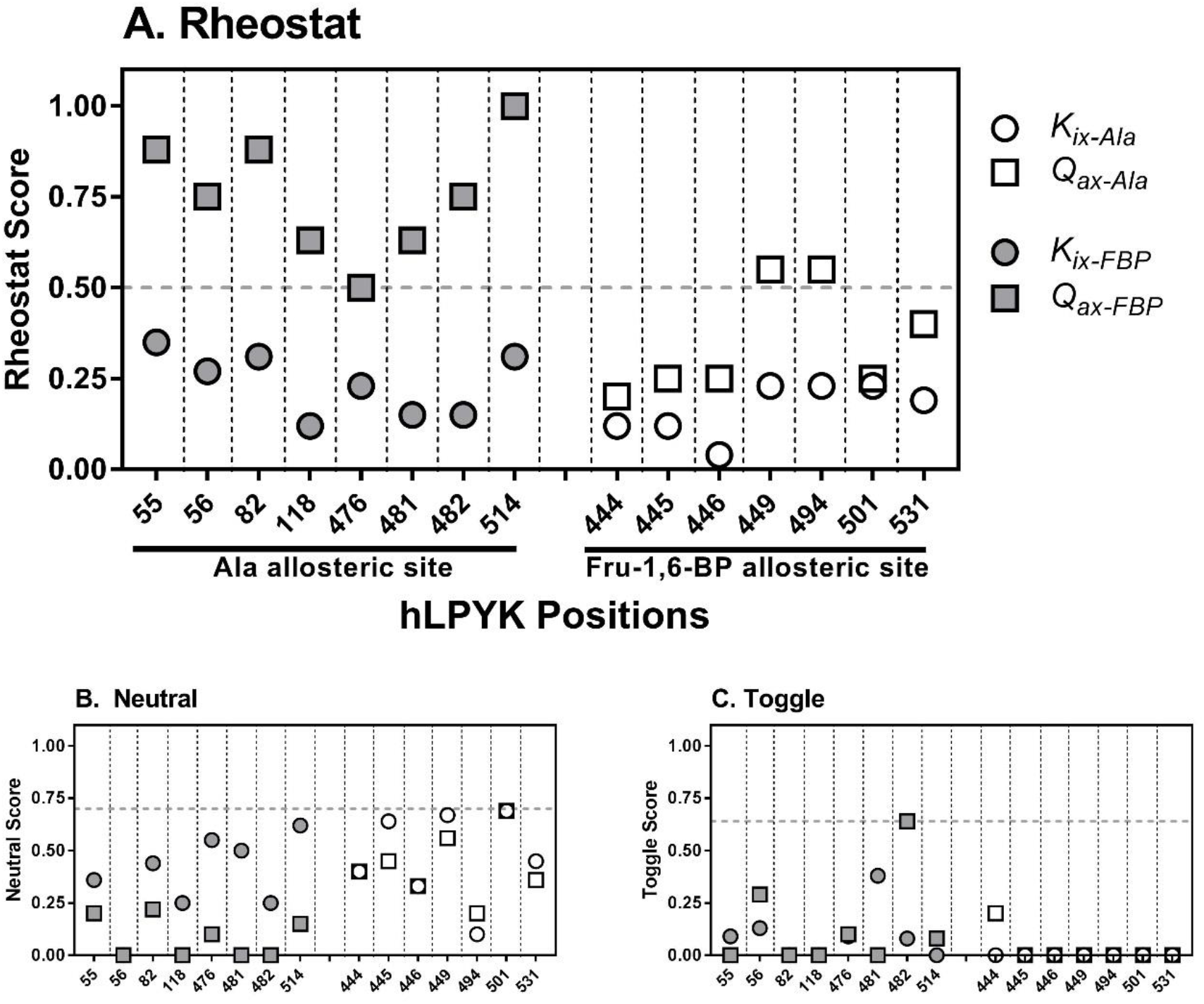
Calculated scores for substituted “distant” positions in and near the allosteric sites of hLPYK. The effects of distant substitutions in the Fru-1,6-BP binding site on Ala binding (*K*_*ix-Ala*_) and PEP/Ala allosteric coupling (*Q*_*ax-Ala*_) and the effects of distant substitutions in the Ala binding site on Fru-1,6-BP binding (*K*_*ix-FBP*_) and PEP/Fru-1,6-BP allosteric coupling (*Q*_*ax-FBP*_) were used to calculate rheostat (top), neutral (bottom left) and toggle (bottom right) scores for the hLPYK positions noted on the x-axis. The scores for the same variants on their respective “local” parameters were previously reported^3^. For direct comparison with the distant data and as described in Methods, the revised scores calculated for the local dataset are included in an all-inclusive figure in the supplement. Dashed vertical lines separate the scores for individual positions. For the rheostat scores, the horizontal dashed line at 0.5 indicates the empirically determined threshold that delineates rheostat positions. For the neutral scores, the horizontal dashed line at 0.7 to delineate high neutral scores is based on empirical comparisons in a previous study ^12^. For the toggle scores, the horizontal dashed line at 0.64 corresponds to the threshold defined in ^14^ to delineate a toggle position.

The right-hand side of Figure 3 shows the influence of positions in the Fru-1,6-BP binding site on Ala binding and PEP/Ala allosteric coupling. For most positions, the rheostat scores were low for both parameters. Nonetheless, position 494 had a rheostat score for *Q*_*ax-Ala*_ above 0.5, indicating distant control over PEP/Ala coupling. Again, the influence on allosteric coupling was often greater than effects on effector binding. Again, none of the parameters demonstrated neutral scores above 0.7. Likewise, there was very little toggle character for positions in the Fru-1,6-BP binding site for Ala binding or PEP/Ala allosteric coupling with no scores above the 0.64 toggle score cutoff.

In summary, substitution of one position in the Fru-1,6-BP binding site significantly altered PEP/Ala coupling and several positions in the Ala binding site exhibited rheostatic control over Fru-1,6-BP allosteric coupling. Strikingly, for *Q*_*ax-FBP*_, position 514 exhibited a nearly perfect rheostat score and positions 55 and 82 had high scores. Indeed, these scores were larger than any identified for the “local” substitution study.

Next, we considered the overall tunability of the two allosteric coupling parameters, *Q*_*ax-Ala*_ and *Q*_*ax-FBP*_. Tunability was determined by combining data for all positions in the Ala site or all positions in the Fru-1,6-BP site (except position 483, as indicated above) for RheoScale analyses. Figure 4 shows full tunability was observed for both parameters: the “better than wildtype” bin and each histogram bin between the maximum (wildtype) and minimum (no allostery) values were occupied by at least one variant. Interestingly, similar results were obtained when the individual position with perfect rheostat control over *Q*_*ax-FBP*_ was excluded from analyses. Thus, the overall tunability of *Q*_*ax-Ala*_ and *Q*_*ax-FBP*_ was very high, even though only a small subset of the total hLPYK positions were used for analyses.

**Figure 4.**
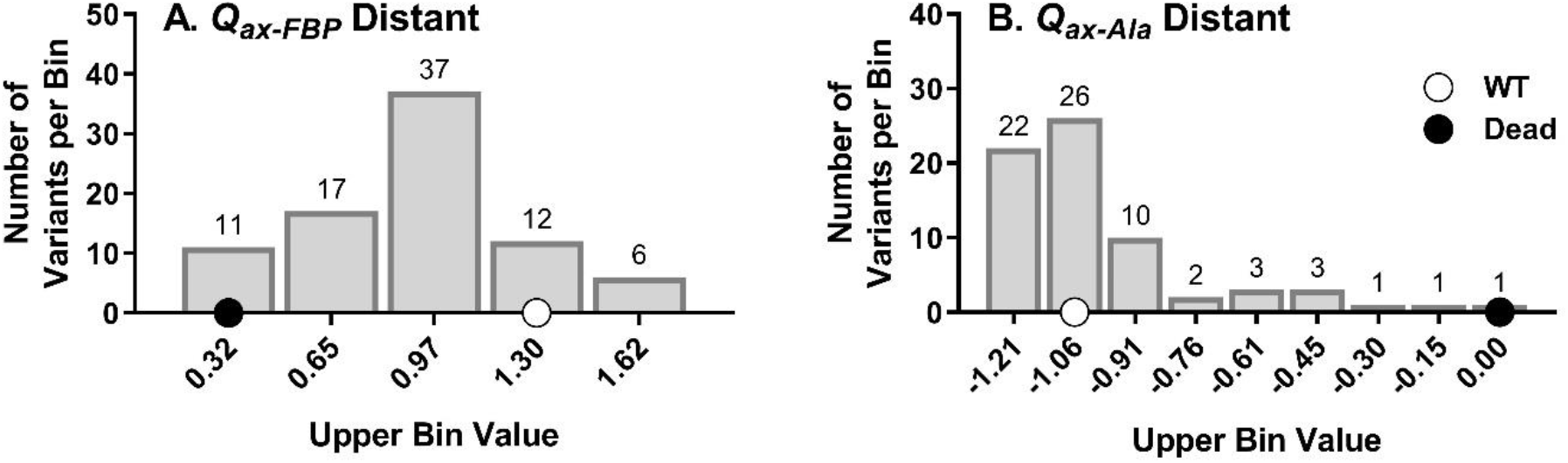
Tunability of allosteric coupling functions *via* substitutions at rheostat positions in “distant” sites. Data shown here are a compilation of the allosteric coupling values measured for all substitutions for all positions in Figure 3. Like Figure 2A, at least one substitution in each binding site results in one of the possible functional outcome subranges (histogram bins) that spans the range between no allostery/”dead” (black dot) and wildtype allosteric coupling (white dot).

Figure 5 shows the rheostat scores calculated for the full dataset, the local subset, and the distant subset. Surprisingly, for *Q*_*ax-FBP*_, the variants located in the Ala binding site (distant) can better tune this parameter (*i.e.*, values closer to 1) than the set of variants located in the Fru-1,6-BP binding site (local). Furthermore, substitutions in the Fru-1,6-BP site (distant) can tune*Q*_*ax-Ala*_ as comprehensively as those in the Ala binding site (local).

**Figure 5.**
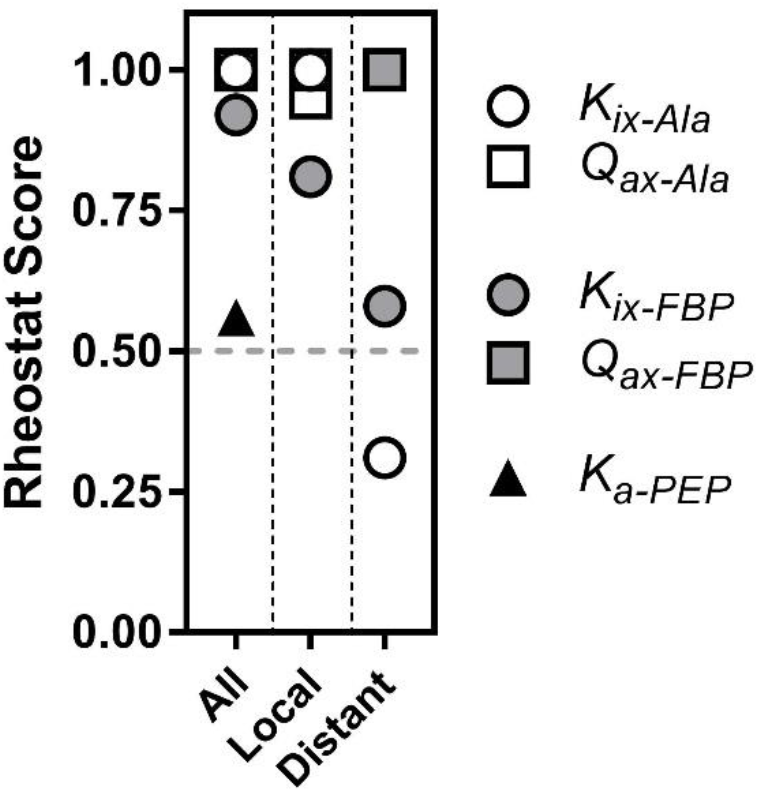
Rheostat scores for compilations of substitutional outcomes from multiple positions. When compilations of data are used to calculate a rheostat score, full tunability is reflected by a rheostat score that approaches 1. Results from both the current work and the previous study^3^ are summarized here: Compilational rheostat scores were calculated for the “local” dataset, the “distant” dataset, and the combination of both by compiling data from the respective sets of positions. When the symbol for *Q*_*ax-Ala*_ is not obvious in the “All” and “Distant” columns, it is behind the symbol for *Q*_*ax-FBP*_.

We next considered additional insights these data might provide about the functional roles of the individual substituted positions. Notably, when both local and distant scores were considered, positions 56, 82, 118, 476, and 514 exerted rheostatic control over multiple parameters (Figure 6). Due to the mathematical relationship of parameters (equation 1), we queried whether correlation existed between rheostatically-controlled parameters with rheostat scores above 0.5 (*e.g.*, Does a tryptophan substitution at position 56 simultaneously change *K*_*ix-Ala*_ and *Q*_*ax-Ala*_?). No correlation was observed (Supplement). Furthermore, when the maximum rheostat and toggle scores were examined for each position (Figure 7), position 482 and 494 were shown to exert rheostatic control over one function and toggle control over a second. Thus, it appears that substitutions at individual hLPYK positions can alter multiple functional parameters.

**Figure 6.**
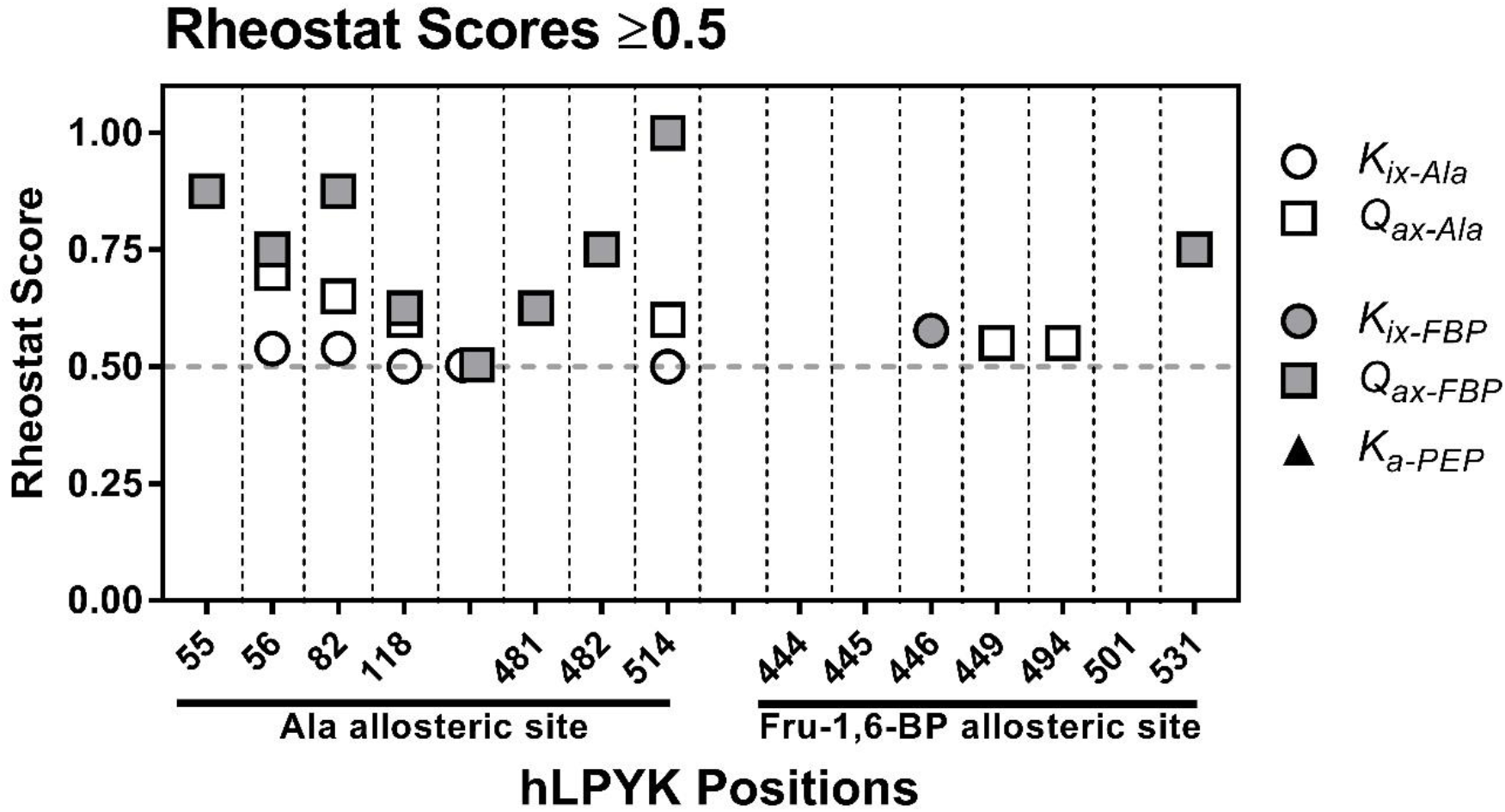
Rheostat scores greater than or equal to 0.5 for both distant and local parameters. Note that no rheostat scores for *K*_*a-PEP*_ met this threshold. Positions 56, 82, 118 and 514 exhibited high rheostat scores for three allosteric parameters and position 476 had high rheostat scores for two parameters.

**Figure 7.**
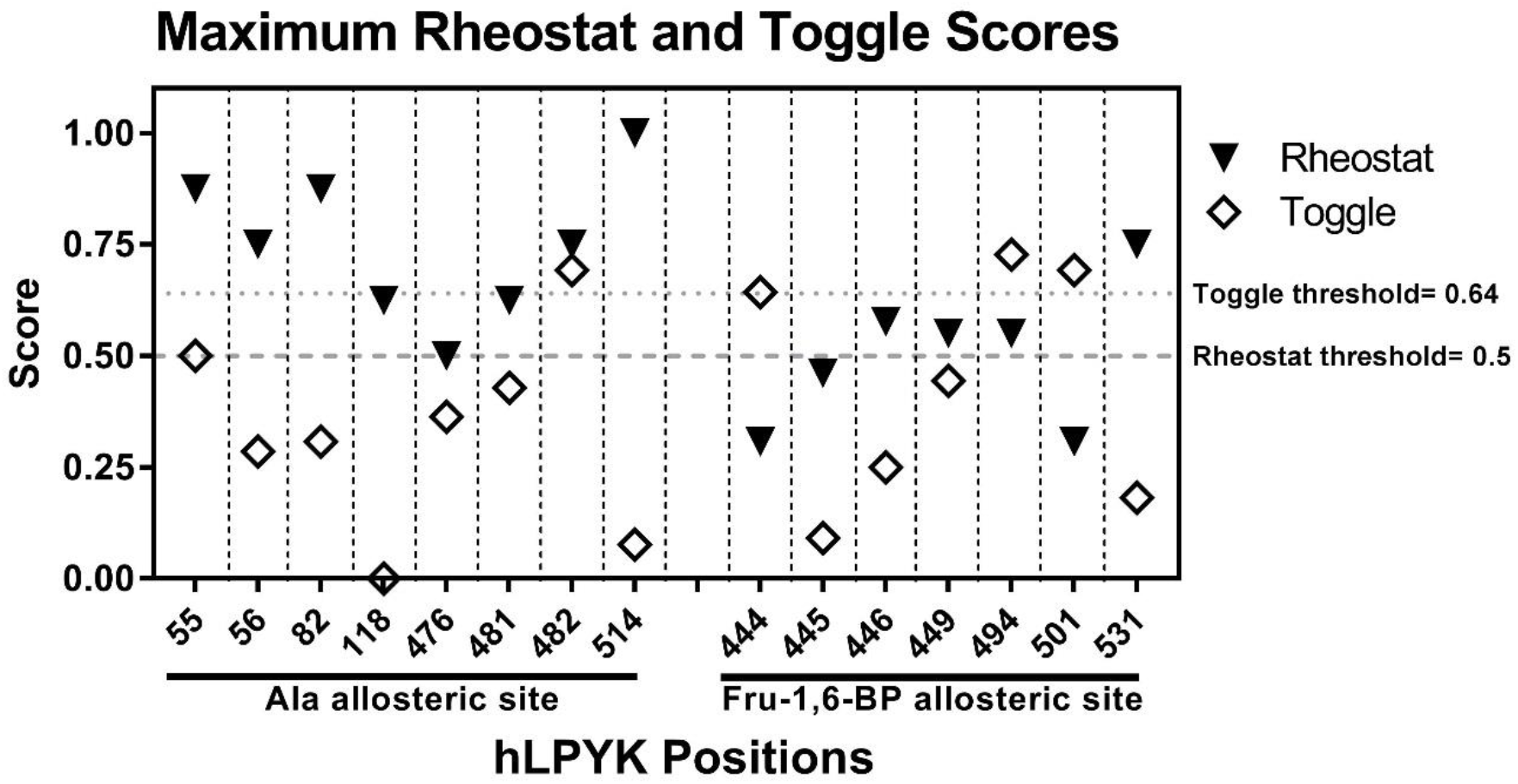
The maximum rheostat and maximum toggle scores for each position when collectively considering all parameters in Table 1. Note that positions 482 and 494 have both a high rheostat score for one functional parameter and a high toggle score for another parameter.

## Discussion

Previously, we identified rheostat positions in the two allosteric sites of hLPYK and showed that allosteric coupling between their respective ligands and substrate binding was fully tunable *via* substitutions at just these subsets of positions. In other words, PEP/Ala allosteric coupling was tunable *via* substitutions in the Ala binding site and PEP/Fru-1,6-BP allosteric coupling was tunable *via* substitutions in the Fru-1,6-BP binding site^3^. Full tunability was indicated by the range of functional changes that resulted from substitutions (Figure 2) that resulted in a rheostat score near 1. The goal of this current study was to determine whether positions distant from the two allosterically-coupled binding sites could also exert rheostatic control over allosteric coupling. To that end, we used the identical variants of the previous study and evaluated their effects on the allosteric parameters for the other binding site.

The results from this study confirm that substitutions distant to the two allosterically-coupled binding sites can exert full tunability over allosteric coupling (*e.g.*, full bin occupancies in Figure 4 and rheostat scores of 1 in Figure 5). Arguably, this study was biased towards success since the chosen “distant” positions were located in sites with known regulatory function. Nonetheless, it is striking that substitutions at these positions have as much or more rheostatic control of the distant parameters as they do of their local parameters (Supplement). Future substitution studies at positions distant to any known allosteric binding sites will be useful to evaluate whether this influence is restricted to allosteric binding sites or common to many regions throughout the protein. We find it possible that rheostatic control of allosteric parameters will be widespread.

The scores shown in Figure 3 do exhibit asymmetry in the distant effects on the allosteric parameters. Substitutions in the Ala binding sites have bigger effects on allosteric parameters than substitutions in the Fru-1,6-BP site. This outcome is consistent with the predictions that several positions in the Ala binding site participate as “choke points” in networks of interactions ^15^. Those predictions are further consistent with outcomes from a full protein alanine scan of hLPYK ^13^.

Another asymmetry is obvious when comparing the distant effects on *Q*_*ax*_ values to those on *K* values. The effects on *Q*_*ax*_ values were much larger. This observation agrees with the emerging view that the positions which contribute to binding can be distinct from those that contribute to allosteric regulation ^5,8,16,17^.

We anticipate that understanding tunability of function *via* substitution will facilitate better ways to rationally engineer allosteric communication in new proteins and to identify potential relationships between allostery and genetic disease in personalized medicine. Indeed, this work can be summarized as a study of the tunability (*via* substitutions at rheostat positions) of tunability (allosteric regulation). As we pursue engineering and disease correlation goals, we must keep in mind two surprising observations from these studies: one position can have rheostatic control over two or more functional parameters and one position can have rheostatic control of one functional parameter and toggle control over a second (Figure 6 and Figure 7). Future efforts must recognize the potential of dual-function positions.

In summary, our attempt to quantitatively evaluate the toggle/rheostat/neutral character of even a limited number of positions in one protein has highlighted that allosteric coupling is highly tunable, even by positions distant from the two allosterically-coupled binding sites. Furthermore, the overlapping roles for one position in controlling multiple functions illuminates the challenges that must be resolved for protein engineering and effective predictions of functional outcomes from variants in patient genomes.

## Acknowledgments

Research in the Fenton laboratory is supported by NIH grant GM115340. We appreciate the contributions from Qingling Tang, who contributed in many ways to this project.

