## Supplemental Data for "Functional tunability from a distance: Rheostat positions influence allosteric coupling between two distant binding sites"

### Supplemental Information

| **Supplemental Table 1: Wildtype averages** | | |
|---|---|---|
| | **average in mM (n=5)** | **Standard deviation of independent evaluations** |
| $K_{a\text{-}PEP}$ (mM) | 0.24 | 0.02 |
| $K_{ix\text{-}ala}$ (mM) | 0.31 | 0.04 |
| $K_{ix\text{-}FBP}$ (mM) | 0.00018 | 0.00008 |
| $Q_{ax\text{-}ala}$ | 0.073 | 0.009 |
| $Q_{ax\text{-}FBP}$ | 14 | 3 |

$K_{ix\text{-}Ala}$ individual histograms

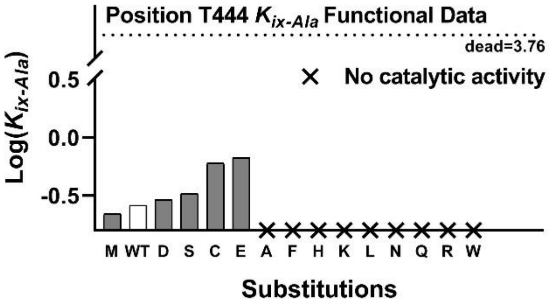

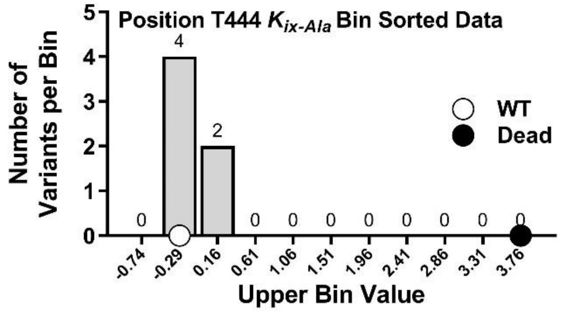

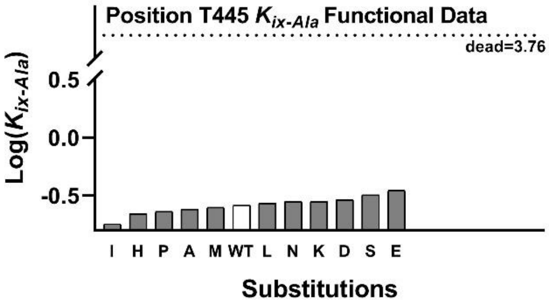

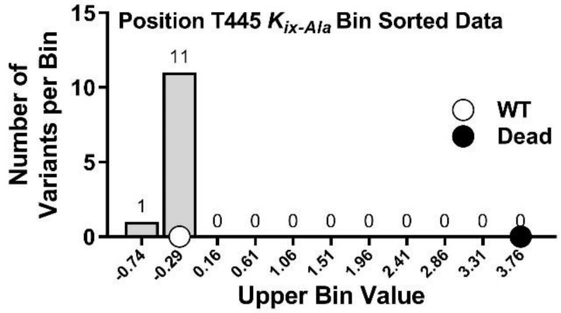

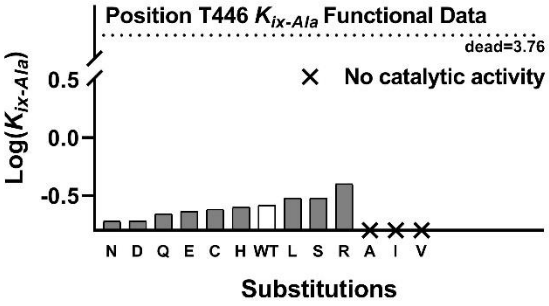

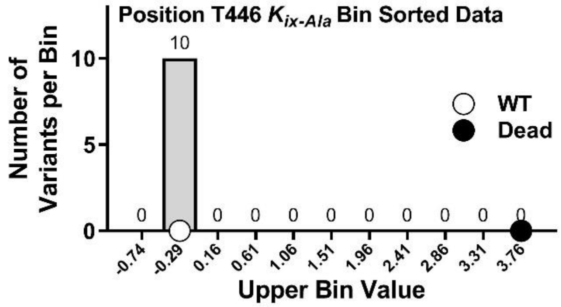

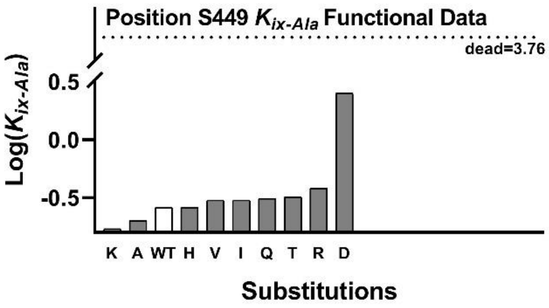

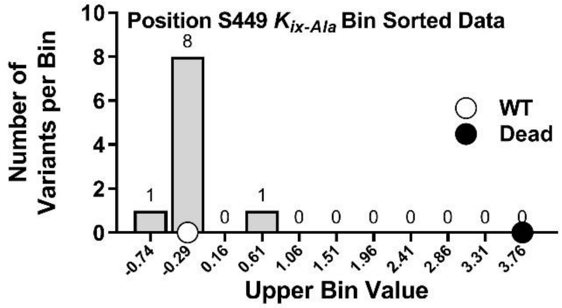

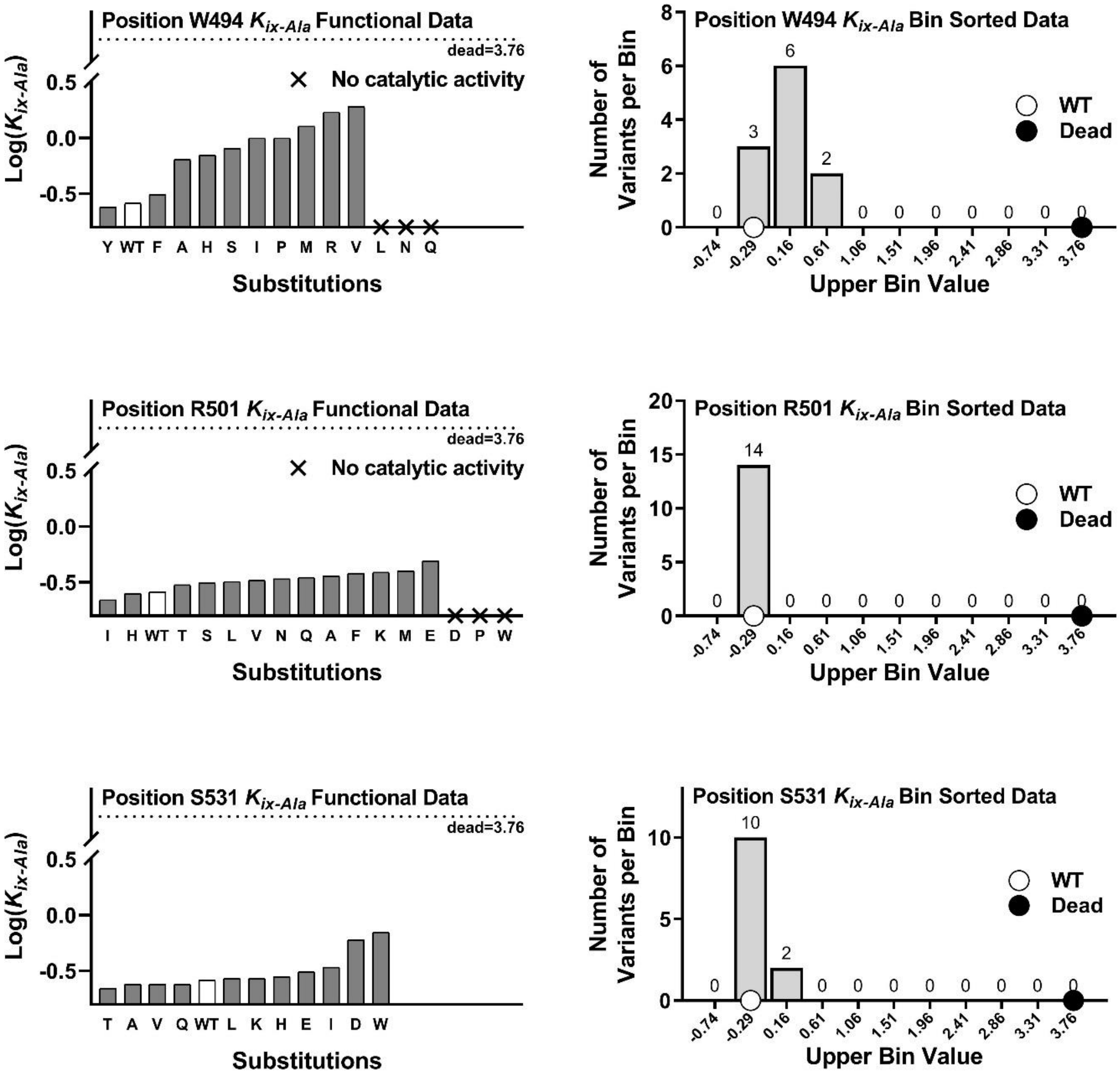
Position W494 $K_{ix-Ala}$ Functional Data
dead=3.76
No catalytic activity
Log($K_{ix-Ala}$)
0.5
0.0
-0.5
Y WT F A H S I P M R V L N Q
Substitutions
Position W494 $K_{ix-Ala}$ Bin Sorted Data
Number of Variants per Bin
8
6
4
2
0
0 3 6 2 0 0 0 0 0 0 0
WT
Dead
-0.74 -0.29 0.16 0.61 1.06 1.51 1.96 2.41 2.86 3.31 3.76
Upper Bin Value
Position R501 $K_{ix-Ala}$ Functional Data
dead=3.76
No catalytic activity
Log($K_{ix-Ala}$)
0.5
0.0
-0.5
I H WT T S L V N Q A F K M E D P W
Substitutions
Position R501 $K_{ix-Ala}$ Bin Sorted Data
Number of Variants per Bin
20
15
10
5
0
0 14 0 0 0 0 0 0 0 0 0
WT
Dead
-0.74 -0.29 0.16 0.61 1.06 1.51 1.96 2.41 2.86 3.31 3.76
Upper Bin Value
Position S531 $K_{ix-Ala}$ Functional Data
dead=3.76
Log($K_{ix-Ala}$)
0.5
0.0
-0.5
T A V Q WT L K H E I D W
Substitutions
Position S531 $K_{ix-Ala}$ Bin Sorted Data
Number of Variants per Bin
10
5
0
0 10 2 0 0 0 0 0 0 0 0
WT
Dead
-0.74 -0.29 0.16 0.61 1.06 1.51 1.96 2.41 2.86 3.31 3.76
Upper Bin Value

*$K_{ix\text{-}FBP}$* individual histograms

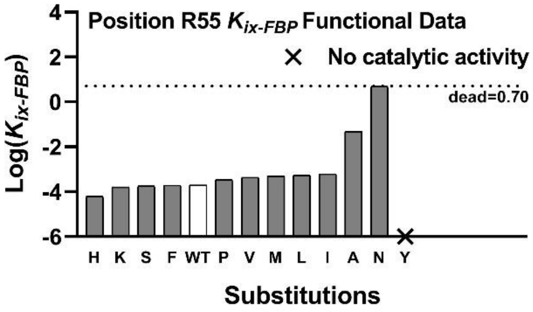

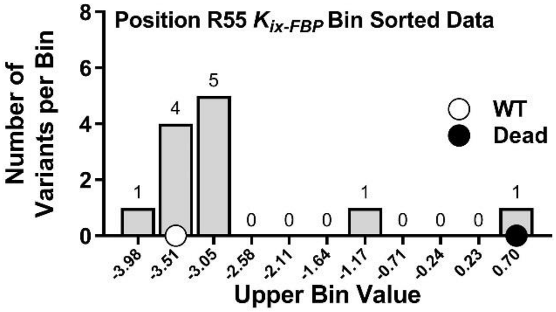

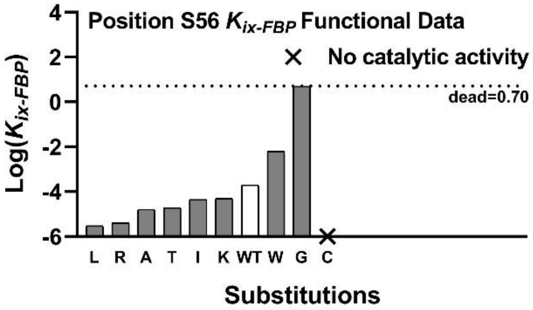

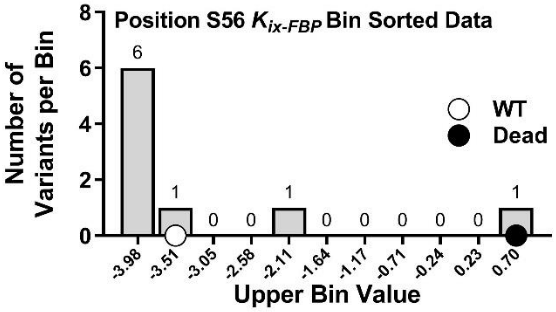

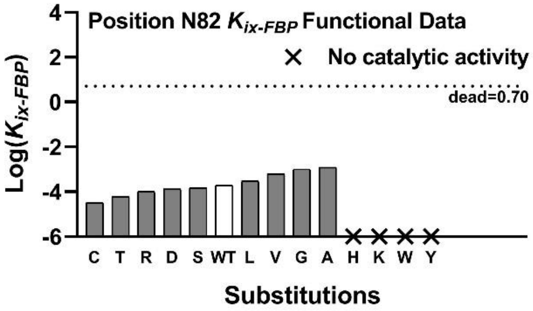

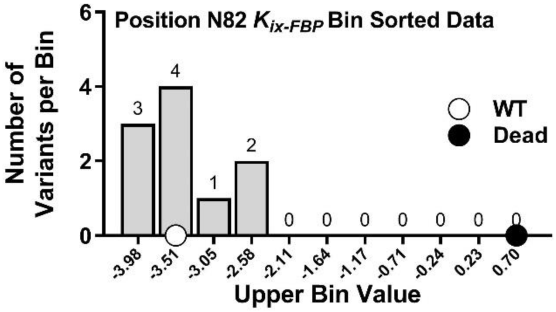

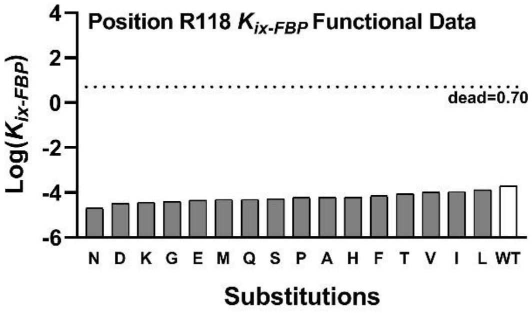

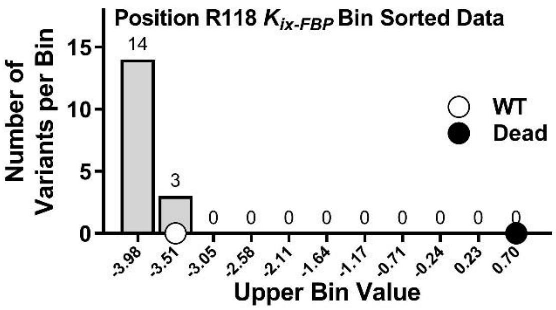

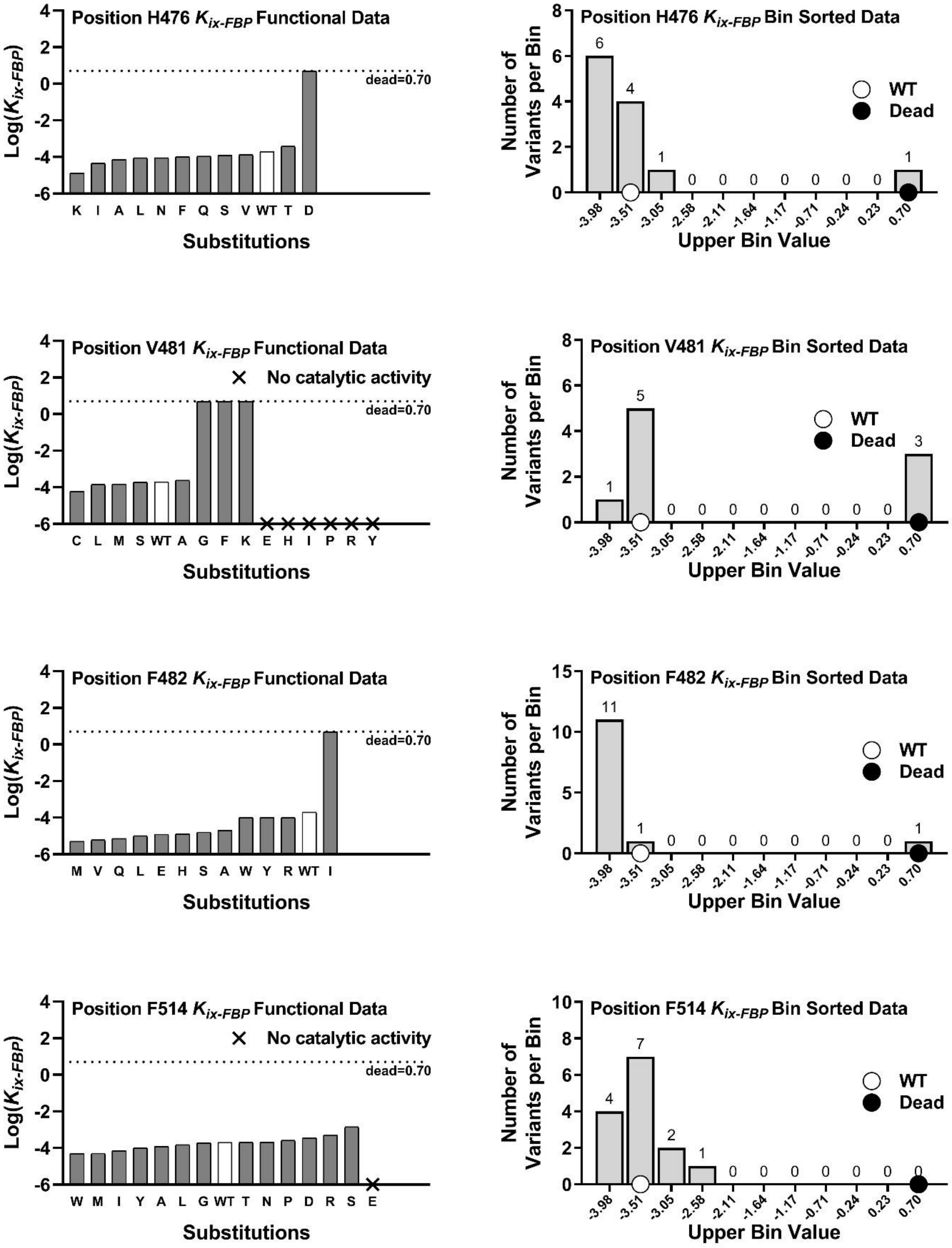

Position H476 $K_{ix-FBP}$ Functional Data
Log($K_{ix-FBP}$)
dead=0.70
K I A L N F Q S V WT T D
Substitutions
Position H476 $K_{ix-FBP}$ Bin Sorted Data
Number of Variants per Bin
WT
Dead
6 4 1 0 0 0 0 0 0 0 1
-3.98 -3.51 -3.05 -2.58 -2.11 -1.64 -1.17 -0.71 -0.24 0.23 0.70
Upper Bin Value
Position V481 $K_{ix-FBP}$ Functional Data
Log($K_{ix-FBP}$)
× No catalytic activity
dead=0.70
C L M S WT A G F K E H I P R Y
Substitutions
Position V481 $K_{ix-FBP}$ Bin Sorted Data
Number of Variants per Bin
WT
Dead
1 5 0 0 0 0 0 0 0 0 3
-3.98 -3.51 -3.05 -2.58 -2.11 -1.64 -1.17 -0.71 -0.24 0.23 0.70
Upper Bin Value
Position F482 $K_{ix-FBP}$ Functional Data
Log($K_{ix-FBP}$)
dead=0.70
M V Q L E H S A W Y R WT I
Substitutions
Position F482 $K_{ix-FBP}$ Bin Sorted Data
Number of Variants per Bin
WT
Dead
11 1 0 0 0 0 0 0 0 0 1
-3.98 -3.51 -3.05 -2.58 -2.11 -1.64 -1.17 -0.71 -0.24 0.23 0.70
Upper Bin Value
Position F514 $K_{ix-FBP}$ Functional Data
Log($K_{ix-FBP}$)
× No catalytic activity
dead=0.70
W M I Y A L G WT T N P D R S E
Substitutions
Position F514 $K_{ix-FBP}$ Bin Sorted Data
Number of Variants per Bin
WT
Dead
4 7 2 1 0 0 0 0 0 0 0
-3.98 -3.51 -3.05 -2.58 -2.11 -1.64 -1.17 -0.71 -0.24 0.23 0.70
Upper Bin Value

$Q_{ax\text{-}Ala}$ individual histograms

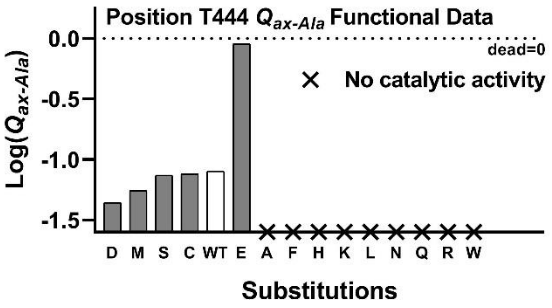

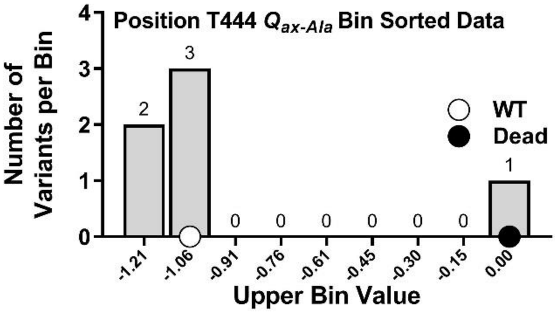

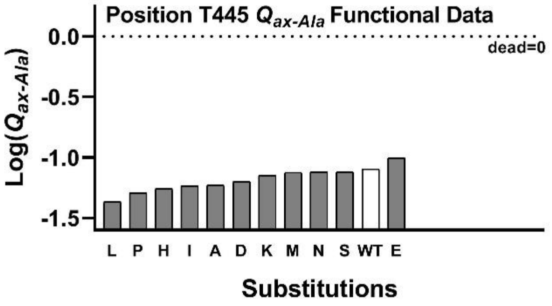

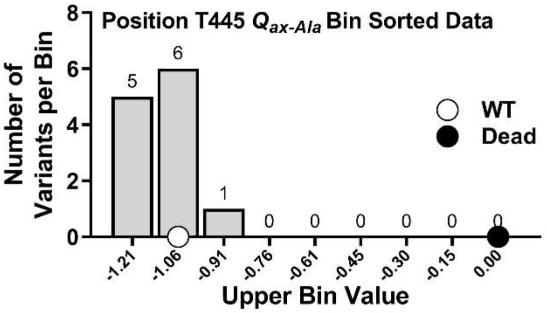

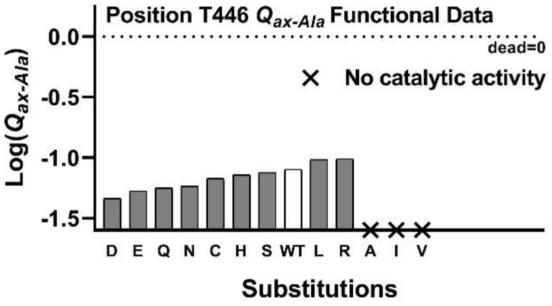

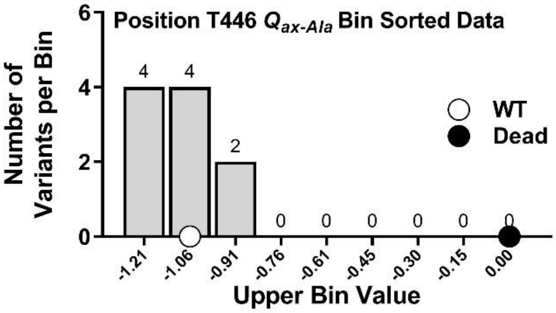

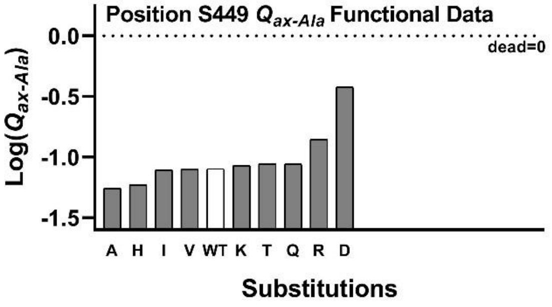

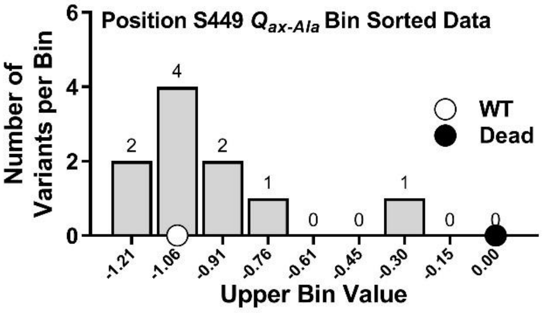

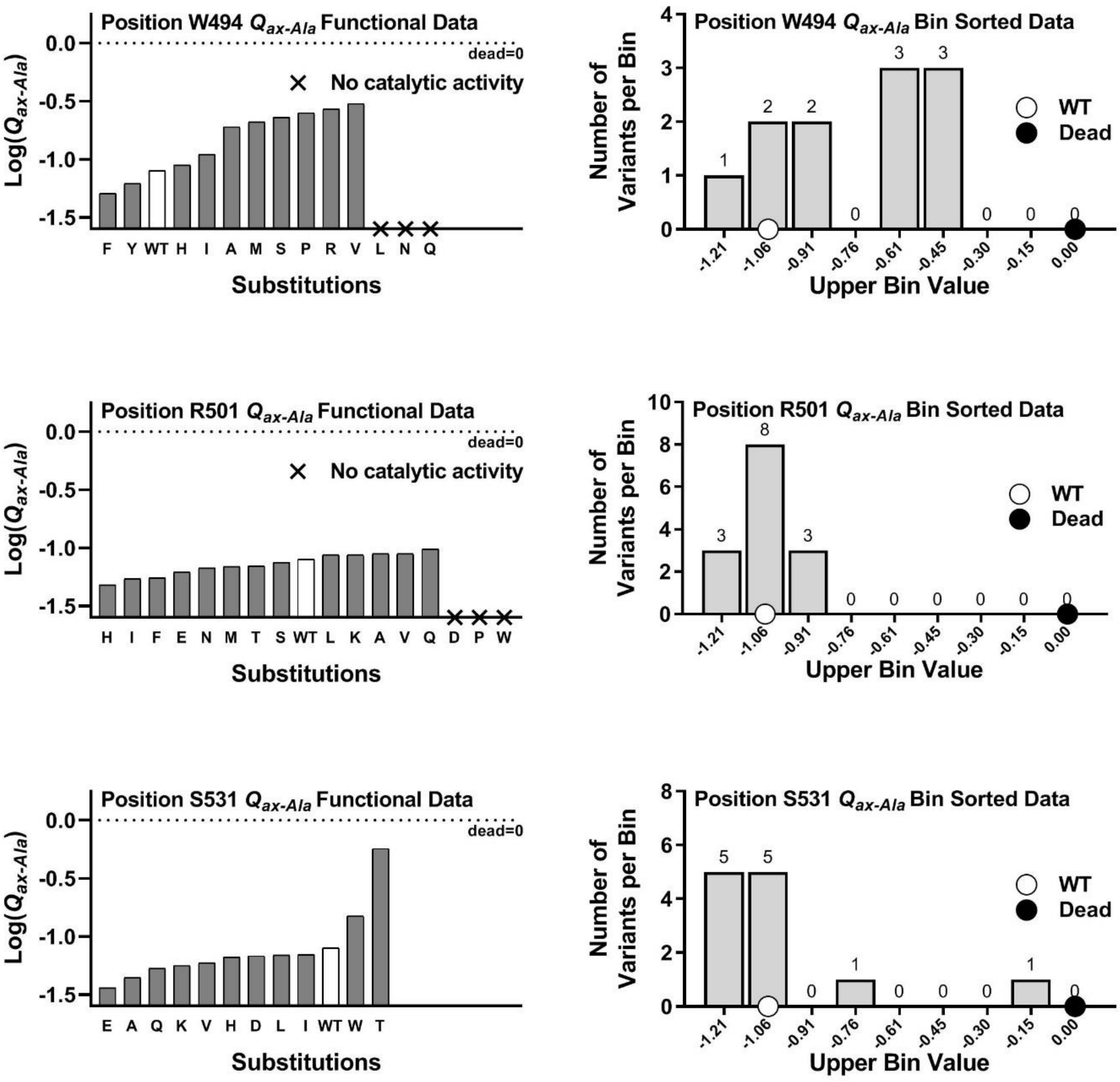
Position W494 $Q_{ax\text{-}Ala}$ Functional Data
Log($Q_{ax\text{-}Ala}$)
0.0
-0.5
-1.0
-1.5
dead=0
× No catalytic activity
F Y WT H I A M S P R V L N Q
Substitutions
Position W494 $Q_{ax\text{-}Ala}$ Bin Sorted Data
Number of Variants per Bin
1 2 2 0 3 3 0 0 0
WT
Dead
-1.21 -1.06 -0.91 -0.76 -0.61 -0.45 -0.30 -0.15 0.00
Upper Bin Value
Position R501 $Q_{ax\text{-}Ala}$ Functional Data
Log($Q_{ax\text{-}Ala}$)
dead=0
× No catalytic activity
H I F E N M T S WT L K A V Q D P W
Substitutions
Position R501 $Q_{ax\text{-}Ala}$ Bin Sorted Data
Number of Variants per Bin
3 8 3 0 0 0 0 0 0
WT
Dead
-1.21 -1.06 -0.91 -0.76 -0.61 -0.45 -0.30 -0.15 0.00
Upper Bin Value
Position S531 $Q_{ax\text{-}Ala}$ Functional Data
Log($Q_{ax\text{-}Ala}$)
dead=0
E A Q K V H D L I WT W T
Substitutions
Position S531 $Q_{ax\text{-}Ala}$ Bin Sorted Data
Number of Variants per Bin
5 5 0 1 0 0 0 1 0
WT
Dead
-1.21 -1.06 -0.91 -0.76 -0.61 -0.45 -0.30 -0.15 0.00
Upper Bin Value

$Q_{ax\text{-}FBP}$ individual histograms

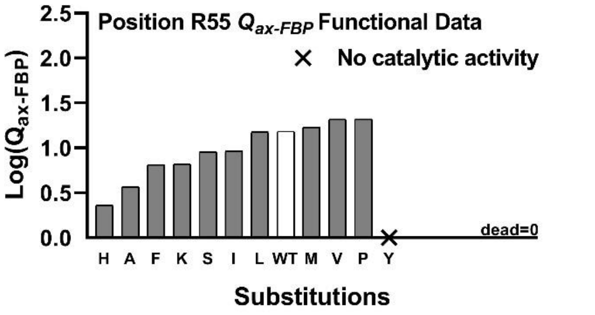

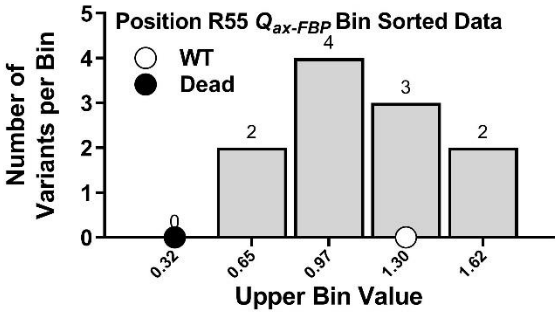

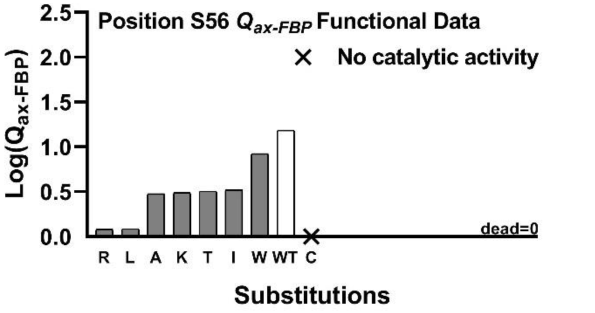

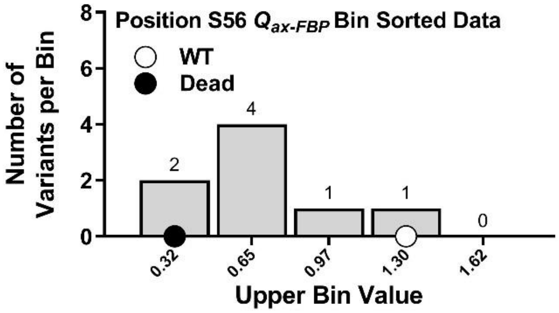

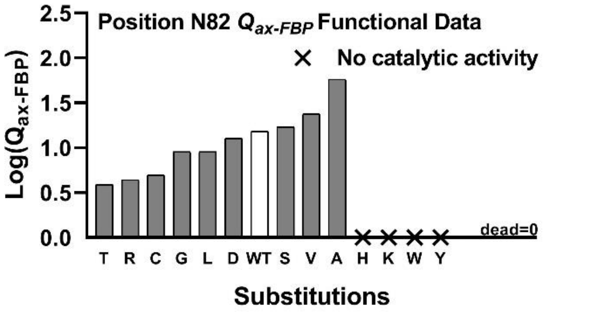

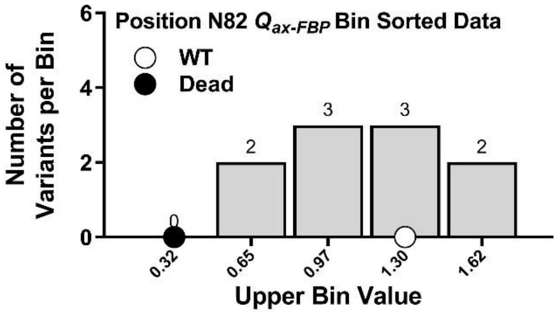

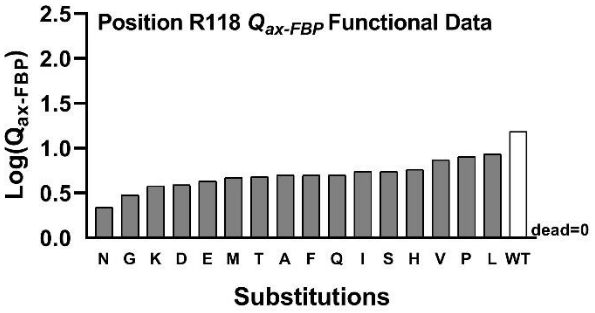

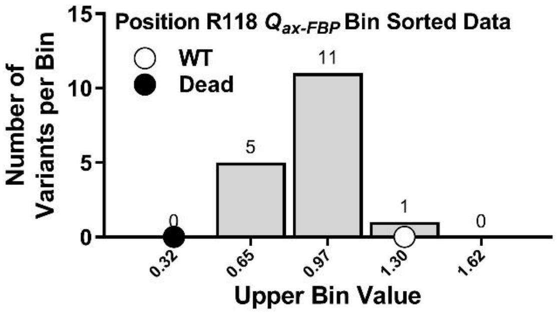

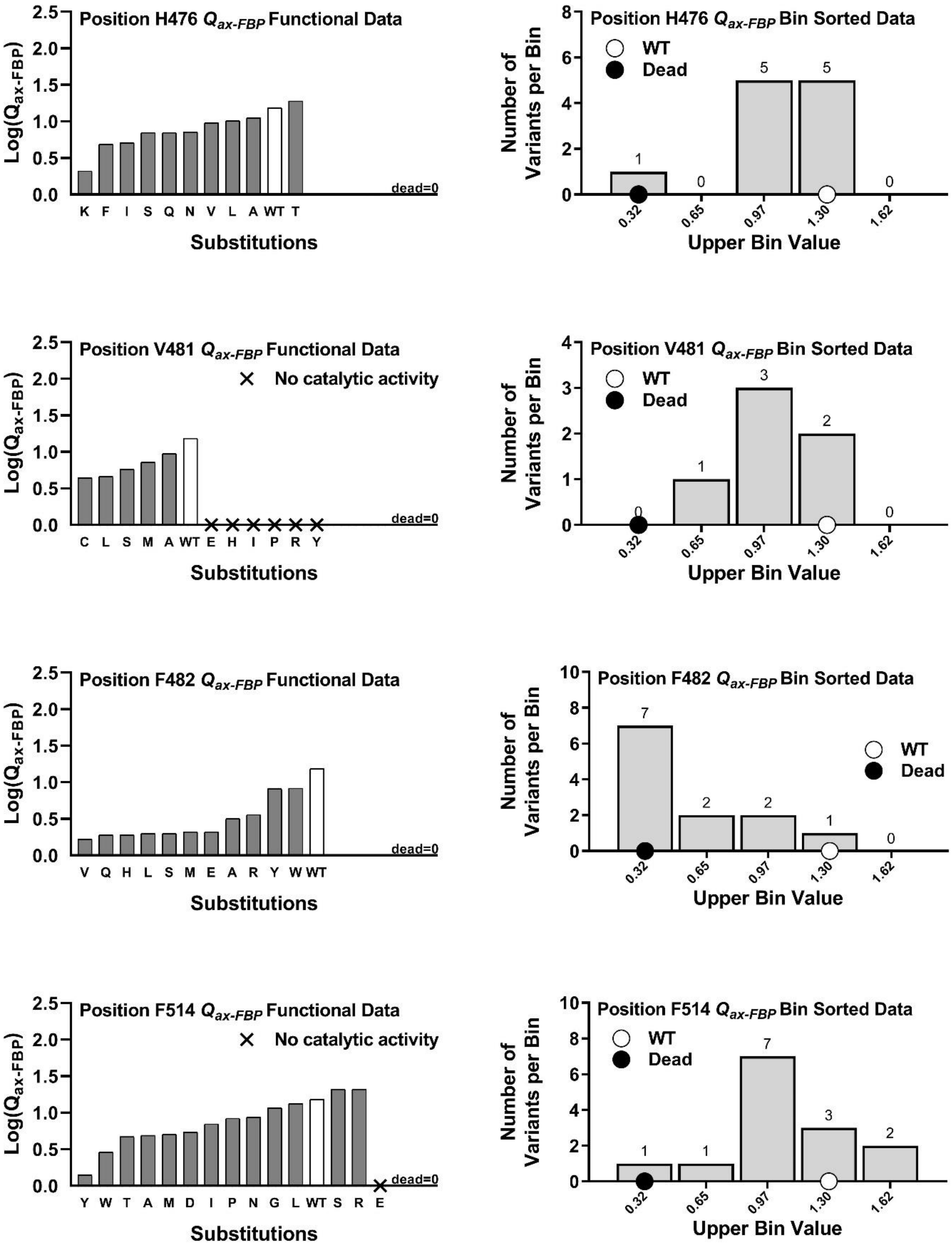

Position H476 $Q_{ax\text{-}FBP}$ Functional Data
Log($Q_{ax\text{-}FBP}$)
2.5
2.0
1.5
1.0
0.5
0.0
dead=0
K F I S Q N V L A WT T
Substitutions
Position H476 $Q_{ax\text{-}FBP}$ Bin Sorted Data
Number of Variants per Bin
WT
Dead
1 0 5 5 0
0.32 0.65 0.97 1.30 1.62
Upper Bin Value
Position V481 $Q_{ax\text{-}FBP}$ Functional Data
× No catalytic activity
Log($Q_{ax\text{-}FBP}$)
dead=0
C L S M A WT E H I P R Y
Substitutions
Position V481 $Q_{ax\text{-}FBP}$ Bin Sorted Data
Number of Variants per Bin
WT
Dead
0 1 3 2 0
0.32 0.65 0.97 1.30 1.62
Upper Bin Value
Position F482 $Q_{ax\text{-}FBP}$ Functional Data
Log($Q_{ax\text{-}FBP}$)
dead=0
V Q H L S M E A R Y W WT
Substitutions
Position F482 $Q_{ax\text{-}FBP}$ Bin Sorted Data
Number of Variants per Bin
WT
Dead
7 2 2 1 0
0.32 0.65 0.97 1.30 1.62
Upper Bin Value
Position F514 $Q_{ax\text{-}FBP}$ Functional Data
× No catalytic activity
Log($Q_{ax\text{-}FBP}$)
dead=0
Y W T A M D I P N G L WT S R E
Substitutions
Position F514 $Q_{ax\text{-}FBP}$ Bin Sorted Data
Number of Variants per Bin
WT
Dead
1 1 7 3 2
0.32 0.65 0.97 1.30 1.62
Upper Bin Value

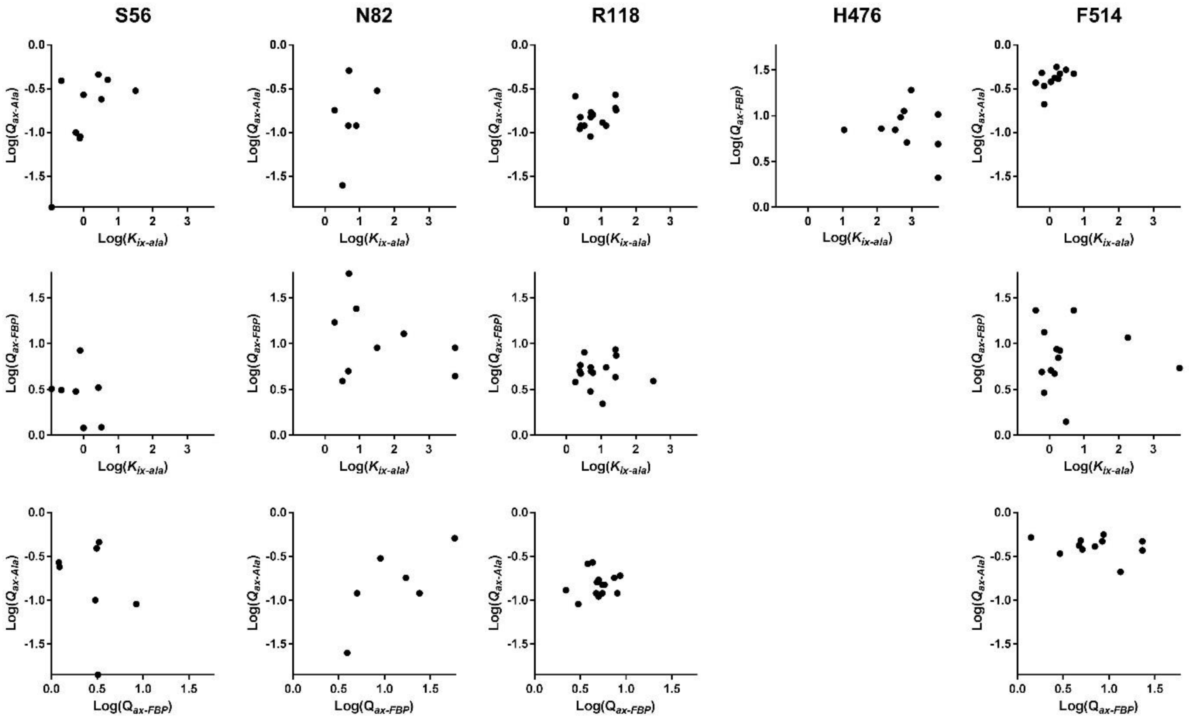

Correlation between parameters. For positions with two or more rheostat scores greater than 0.5, the values of the relevant $K_{a\text{-}PEP}$, $K_{ix\text{-}Ala}$, $K_{ix\text{-}FBP}$, $Q_{ax\text{-}Ala}$, or $Q_{ax\text{-}FBP}$ were compared for each variant (individual dots). No correlation is observed among the parameters, which indicates that each substitution has independent effects on the different functional parameters.

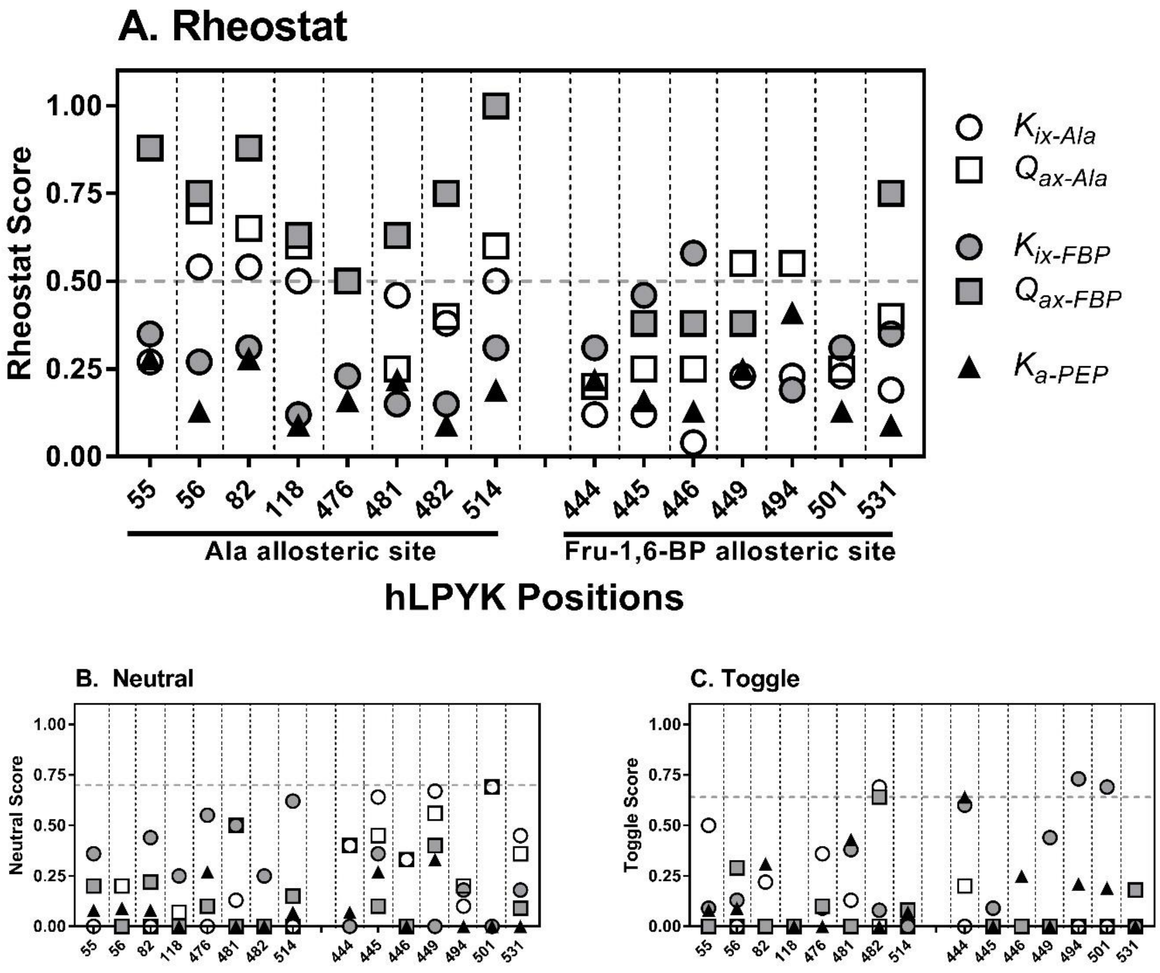

All scores for local and distant evaluations combined.

Supplemental Table 2

| Protein | $K_{a\text{-}PEP}$ [mM] | $K_{ix\text{-}Ala}$ [mM] | $Q_{ax\text{-}Ala}$ |
|---|---|---|---|
| Wildtype[b] | 0.23±0.01 | 0.26±0.01 | 0.080±0.001 |
| **Thr444** | | | |
| T444A | No activity | No activity | No activity |
| T444L | No activity | No activity | No activity |
| T444M | 0.38±0.01 | 0.22±0.02 | 0.055±0.003 |
| T444F | No activity | No activity | No activity |
| T444W | No activity | No activity | No activity |
| T444S | 0.25±0.01 | 0.33±0.03 | 0.074±0.003 |
| T444C | 0.70±0.06 | 0.6±0.1 | 0.076±0.007 |
| T444N | No activity | No activity | No activity |
| T444Q | No activity | No activity | No activity |
| T444H | No activity | No activity | No activity |
| T444K | No activity | No activity | No activity |
| T444R | No activity | No activity | No activity |
| T444D | 0.40±0.01 | 0.29±0.03 | 0.044±0.002 |
| T444E | 0.66±0.02 | 0.67±0.02 | 0.9±0.1 |
| **Thr445** | | | |
| T445A | 0.16±0.01 | 0.24±0.01 | 0.059±0.002 |
| T445L | 0.24±0.01 | 0.27±0.01 | 0.043±0.001 |
| T445I | 0.15±0.01 | 0.18±0.01 | 0.058±0.002 |
| T445M | 0.21±0.01 | 0.25±0.01 | 0.075±0.002 |
| T445P | 0.44±0.01 | 0.23±0.02 | 0.051±0.007 |
| T445S | 0.20±0.01 | 0.32±0.02 | 0.076±0.003 |
| T445N | 0.21±0.01 | 0.28±0.01 | 0.076±0.001 |
| T445H | 0.20±0.01 | 0.22±0.01 | 0.055±0.001 |
| T445K | 0.19±0.01 | 0.28±0.01 | 0.071±0.002 |
| T445D | 0.22±0.01 | 0.29±0.01 | 0.063±0.002 |
| T445E | 0.26±0.01 | 0.35±0.01 | 0.099±0.001 |
| **Thr446** | | | |
| T446A | No activity | No activity | No activity |
| T446V | No activity | No activity | No activity |
| T446L | 0.11±0.01 | 0.30±0.02 | 0.096±0.004 |
| T446I | No activity | No activity | No activity |
| T446S | 0.20±0.01 | 0.30±0.01 | 0.075±0.002 |
| T446C | 0.20±0.01 | 0.24±0.02 | 0.067±0.004 |
| T446N | 0.15±0.01 | 0.19±0.01 | 0.058±0.002 |
| T446Q | 0.12±0.01 | 0.22±0.05 | 0.056±0.006 |
| T446H | 0.20±0.01 | 0.25±0.01 | 0.072±0.001 |
| T446R | 0.20±0.01 | 0.40±0.02 | 0.098±0.003 |

| | | | |
|---|---|---|---|
| T446D | 0.21±0.01 | 0.19±0.01 | 0.046±0.002 |
| T446E | 0.20±0.01 | 0.23±0.02 | 0.053±0.002 |
| **Ser449** | | | |
| S449A | 0.25±0.01 | 0.20±0.01 | 0.055±0.001 |
| S449V | 0.32±0.01 | 0.30±0.02 | 0.079±0.003 |
| S449I | 0.23±0.01 | 0.30±0.03 | 0.078±0.004 |
| S449T | 0.18±0.01 | 0.32±0.03 | 0.088±0.004 |
| S449Q | 0.25±0.01 | 0.31±0.02 | 0.088±0.003 |
| S449H | 0.17±0.01 | 0.26±0.02 | 0.059±0.002 |
| S449K | 0.12±0.01 | 0.17±0.01 | 0.085±0.004 |
| S449R | 0.43±0.01 | 0.38±0.03 | 0.14±0.01 |
| S449D | 1.2±0.1 | 2.5±0.3 | 0.38±0.01 |
| **Trp494** | | | |
| W494A | 0.91±0.02 | 0.64±0.07 | 0.19±0.01 |
| W494V | 1.71±0.05 | 1.9±0.3 | 0.30±0.01 |
| W494L | No activity | No activity | No activity |
| W494I | 0.89±0.05 | 1.0±0.2 | 0.11±0.01 |
| W494M | 1.29±0.01 | 1.27±0.05 | 0.21±0.01 |
| W494P | 1.42±0.03 | 1.0±0.1 | 0.25±0.01 |
| W494F | 0.10±0.01 | 0.31±0.03 | 0.051±0.002 |
| W494S | 1.27±0.07 | 0.8±0.2 | 0.23±0.01 |
| W494Y | 0.14±0.01 | 0.24±0.01 | 0.062±0.002 |
| W494N | No activity | No activity | No activity |
| W494Q | No activity | No activity | No activity |
| W494H | 0.096±0.007 | 0.7±0.1 | 0.089±0.007 |
| W494R | 1.60±0.05 | 1.7±0.2 | 0.27±0.01 |
| **Arg501** | | | |
| R501A | 0.14±0.01 | 0.36±0.01 | 0.089±0.001 |
| R501V | 0.13±0.01 | 0.33±0.04 | 0.089±0.004 |
| R501L | 0.11±0.03 | 0.32±0.02 | 0.087±0.003 |
| R501I | 0.092±0.002 | 0.22±0.01 | 0.054±0.001 |
| R501M | 0.15±0.01 | 0.40±0.02 | 0.069±0.002 |
| R501F | 0.051±0.002 | 0.38±0.03 | 0.055±0.003 |
| R501W | No activity | No activity | No activity |
| R501S | 0.091±0.003 | 0.31±0.03 | 0.075±0.002 |
| R501T | 0.063±0.002 | 0.30±0.02 | 0.070±0.003 |
| R501N | 0.065±0.006 | 0.34±0.07 | 0.067±0.008 |
| R501Q | 0.17±0.01 | 0.35±0.01 | 0.098±0.002 |
| R501H | 0.051±0.002 | 0.25±0.03 | 0.048±0.002 |
| R501K | 0.20±0.01 | 0.39±0.01 | 0.087±0.002 |
| R501D | No activity | No activity | No activity |
| R501E | 0.12±0.01 | 0.49±0.03 | 0.062±0.002 |
| **Ser531** | | | |
| S531A | 0.096±0.003 | 0.24±0.01 | 0.044±0.002 |
| S531V | 0.11±0.01 | 0.24±0.03 | 0.059±0.004 |

| | | | |
|---|---|---|---|
| S531L | 0.13±0.01 | 0.27±0.02 | 0.069±0.002 |
| S531I | 0.11±0.01 | 0.34±0.03 | 0.070±0.003 |
| S531W | 0.32±0.03 | 0.70±0.13 | 0.15±0.01 |
| S531T | 0.17±0.01 | 0.22±0.01 | 0.57±0.002 |
| S531Q | 0.12±0.01 | 0.24±0.01 | 0.053±0.001 |
| S531H | 0.14±0.01 | 0.28±0.01 | 0.066±0.002 |
| S531K | 0.12±0.01 | 0.27±0.01 | 0.056±0.002 |
| S531D | 0.045±0.002 | 0.60±0.05 | 0.068±0.003 |
| S531E | 0.037±0.002 | 0.31±0.02 | 0.036±0.002 |

Supplemental Table 3

| Protein | $K_{a\text{-}PEP}$ [mM] | $K_{ix\text{-}FBP}$ [mM] | $Q_{ax\text{-}FBP}$ |
|---|---|---|---|
| Wildtype | 0.250±0.003 | 0.000197±0.000001 | 15.3±0.3 |
| **Arg55** | | | |
| R55A[b] | 0.056±0.001 | 0.048±0.004 | 3.70±0.09 |
| R55V | 0.52±0.01 | 0.00043±0.00007 | 24±2 |
| R55L | 0.43±0.01 | 0.00055±0.000005 | 15±1 |
| R55I | 0.42±0.02 | 0.0006±0.0002 | 9.3±0.9 |
| R55M | 0.31±0.02 | 0.0005±0.0001 | 17±1 |
| R55P | 0.60±0.02 | 0.00035±0.00004 | 26±1 |
| R55F | 0.15±0.01 | 0.00019±0.00002 | 6.5±0.2 |
| R55S | 0.21±0.01 | 0.00018±0.00004 | 9.0±0.6 |
| R55N | 0.036±0.003 | No activation | No activation |
| R55Y | No activity | No activity | No activity |
| R55H | 0.062±0.001 | 0.00006±0.00001 | 2.3±0.1 |
| R55K | 0.14±0.01 | 0.00016±0.00007 | 6.6±0.9 |
| **Ser56** | | | |
| S56G | 0.026±0.001 | No activation | No activation |
| S56A | 0.079±0.005 | 0.000016±0.000005 | 3.0±0.2 |
| S56L | 0.026±0.001 | 0.000003±0.000004 | 1.22±0.06 |
| S56I | 0.093±0.003 | 0.000043±0.000008 | 3.3±0.1 |
| S56W | 0.29±0.01 | 0.006±0.001 | 8.4±0.9 |
| S56T | 0.069±0.002 | 0.000019±0.000003 | 3.2±0.1 |
| S56C | No activity | No activity | No activity |
| S56K | 0.082±0.004 | 0.00005±0.00002 | 3.1±0.2 |
| S56R | 0.022±0.001 | 0.000004±0.000005 | 1.20±0.07 |
| **Asn82** | | | |
| N82G | 0.26±0.01 | 0.0010±0.0004 | 9±2 |
| N82A | 0.99±0.01 | 0.0012±0.0001 | 58±2 |
| N82V | 0.53±0.02 | 0.0006±0.0001 | 24±3 |
| N82L | 0.26±0.01 | 0.0003±0.0001 | 9±1 |
| N82W | No activity | No activity | No activity |
| N82S | 0.35±0.01 | 0.00015±0.00003 | 17±1 |
| N82T | 0.082±0.007 | 0.00006±0.00002 | 3.9±0.4 |
| N82C | 0.14±0.01 | 0.000032±0.000005 | 5.0±0.2 |
| N82Y | No activity | No activity | No activity |
| N82H | No activity | No activity | No activity |
| N82K | No activity | No activity | No activity |
| N82R | 0.12±0.01 | 0.00010±0.00002 | 4.4±0.2 |
| N82D | 0.42±0.01 | 0.00013±0.00002 | 12.8±0.5 |

| | | | |
|---|---|---|---|
| **Arg118** | | | |
| R118G | 0.080±0.003 | 0.00004±0.00001 | 3.0±0.2 |
| R118A | 0.15±0.01 | 0.000062±0.000007 | 5.0±0.3 |
| R118V | 0.16±0.01 | 0.00010±0.00002 | 7.4±0.4 |
| R118L | 0.17±0.01 | 0.00013±0.00004 | 8.6±0.6 |
| R118I | 0.13±0.01 | 0.00011±0.00002 | 5.5±0.2 |
| R118M | 0.12±0.01 | 0.00005±0.00001 | 4.7±0.2 |
| R118P | 0.16±0.01 | 0.00006±0.00001 | 8.0±0.4 |
| R118F | 0.11±0.01 | 0.00007±0.00001 | 5.0±0.4 |
| R118S | 0.14±0.01 | 0.000052±0.000004 | 5.5±0.2 |
| R118T | 0.12±0.01 | 0.00009±0.00001 | 4.8±0.3 |
| R118N | 0.066±0.006 | 0.00002±0.00001 | 2.2±0.2 |
| R118Q | 0.13±0.01 | 0.00005±0.00001 | 5.0±0.3 |
| R118H | 0.14±0.01 | 0.000062±0.000008 | 5.8±0.2 |
| R118K | 0.099±0.004 | 0.000036±0.000008 | 3.8±0.2 |
| R118D | 0.11±0.01 | 0.000033±0.000004 | 3.9±0.1 |
| R118E | 0.12±0.01 | 0.000045±0.000009 | 4.3±0.2 |
| **His476** | | | |
| H476A | 0.27±0.01 | 0.000069±0.000007 | 11.2±0.03 |
| H476V | 0.18±0.01 | 0.00013±0.00002 | 9.6±0.9 |
| H476L | 0.22±0.01 | 0.00009±0.00001 | 10.3±0.4 |
| H476I | 0.11±0.01 | 0.000046±0.000006 | 5.1±0.2 |
| H476F | 0.12±0.01 | 0.00010±0.00002 | 4.9±0.2 |
| H476S | 0.23±0.01 | 0.00012±0.00003 | 7±1 |
| H476T | 0.44±0.01 | 0.00038±0.00003 | 19±1 |
| H476N | 0.20±0.01 | 0.00009±0.00002 | 7.2±0.4 |
| H476Q | 0.16±0.01 | 0.00011±0.00002 | 7.0±0.04 |
| H476K | 0.056±0.001 | 0.000013±0.000005 | 2.1±0.1 |
| H476D | 0.027±0.002 | No activation | No activation |
| **Val481** | | | |
| V481G | 0.024±0.001 | No activation | No activation |
| V481A | 0.21±0.01 | 0.00024±0.00003 | 9.5±0.7 |
| V481L | 0.12±0.01 | 0.00014±0.00008 | 4.6±0.7 |
| V481I | No activity | No activity | No activity |
| V481M | 0.24±0.02 | 0.00015±0.00005 | 7.2±0.8 |
| V481P | No activity | No activity | No activity |
| V481F | 0.76±0.028 | No activation | No activation |
| V481S | 0.13±0.01 | 0.00019±0.00006 | 5.8±.07 |
| V481C | 0.092±0.004 | 0.00006±0.00001 | 4.4±0.2 |
| V481Y | No activity | No activity | No activity |
| V481H | No activity | No activity | No activity |
| V481K | 0.16±0.01 | No activation | No activation |
| V481R | No activity | No activity | No activity |
| V481E | No activity | No activity | No activity |
| **Phe482** | | | |
| F482A | 0.069±0.003 | 0.000020±0.000005 | 3.2±0.2 |

| | | | |
|---|---|---|---|
| F482V | 0.033±0.001 | 0.000006±0.000004 | 1.66±0.08 |
| F482L | 0.045±0.002 | 0.000010±0.000004 | 2.0±0.1 |
| F482I | 0.032±0.002 | No activation | No activation |
| F482M | 0.038±0.002 | 0.000005±0.000002 | 2.1±0.1 |
| F482W | 0.21±0.01 | 0.00010±0.00001 | 8.3±0.4 |
| F482S | 0.041±0.001 | 0.000016±0.000005 | 2.0±0.1 |
| F482Q | 0.039±0.002 | 0.000007±0.000003 | 1.9±0.1 |
| F482Y | 0.20±0.01 | 0.00010±0.00002 | 8.1±0.5 |
| F482H | 0.048±0.003 | 0.000013±0.000005 | 1.9±0.1 |
| F482R | 0.076±0.0004 | 0.00010±0.00003 | 3.6±0.3 |
| F482E | 0.044±0.002 | 0.000012±0.000005 | 2.1±0.1 |
| **Pro483** | | | |
| P483G | 0.13±0.01 | 0.000045±0.000005 | 4.9±0.2 |
| P483A | No activity | No activity | No activity |
| P483V | No activity | No activity | No activity |
| P483L | No activity | No activity | No activity |
| P483I | No activity | No activity | No activity |
| P483W | No activity | No activity | No activity |
| P483T | No activity | No activity | No activity |
| P483C | No activity | No activity | No activity |
| P483K | No activity | No activity | No activity |
| P483D | No activity | No activity | No activity |
| P483E | No activity | No activity | No activity |
| **Phe514** | | | |
| F514G | 0.27±0.01 | 0.00019±0.00002 | 11.6±0.5 |
| F514A | 0.16±0.01 | 0.00012±0.00004 | 4.9±0.5 |
| F514L | 0.28±0.01 | 0.00015±0.00001 | 13.3±0.4 |
| F514I | 0.17±0.01 | 0.00007±0.00001 | 7.0±0.3 |
| F514M | 0.13±0.01 | 0.000051±0.000006 | 5.1±0.2 |
| F514P | 0.14±0.01 | 0.00026±0.00006 | 8.4±0.7 |
| F514W | 0.078±0.005 | 0.000049±0.000008 | 2.9±0.3 |
| F514S | 0.36±0.02 | 0.0014±0.0004 | 23±3 |
| F514T | 0.14±0.01 | 0.00020±0.00005 | 4.7±0.5 |
| F514N | 0.18±0.01 | 0.00020±0.00003 | 8.7±0.4 |
| F514Y | 0.024±0.01 | 0.0001±0.0001 | 1.4±0.1 |
| F514R | 0.40±0.03 | 0.0005±0.0001 | 23±3 |
| F514D | 0.094±0.002 | 0.00035±0.00008 | 5.4±0.4 |
| F514E | No activity | No activity | No activity |
